# Antibiotic skeletal diversification via differential enoylreductase recruitment and module iteration in *trans*-acyltransferase polyketide synthases

**DOI:** 10.1101/2023.11.30.569433

**Authors:** Xinyun Jian, Fang Pang, Christian Hobson, Matthew Jenner, Lona M. Alkhalaf, Gregory L. Challis

## Abstract

Microorganisms are remarkable chemists capable of assembling complex molecular architectures that penetrate cells and bind biomolecular targets with exquisite selectivity. Consequently, microbial natural products have wide-ranging applications in medicine and agriculture. How the “blind watchmaker” of evolution creates skeletal diversity is a key question in contemporary natural products research. Comparative analysis of biosynthetic pathways to structurally related metabolites is an insightful approach to addressing this.

Here we report comparative biosynthetic investigations of gladiolin, a polyketide antibiotic from *Burkholderia gladioli* with promising activity against multidrug resistant *Mycobacterium tuberculosis*, and entangien, a structurally related antibiotic produced by *Sorangium cellulosum*. Although these metabolites have very similar macrolide cores, their C21 side chains differ significantly in both length and degree of saturation. Surprisingly, the *trans*-acyltransferase polyketide synthases (PKSs) that assemble these antibiotics are almost identical, raising intriguing questions about mechanisms underlying structural diversification in this important class of biosynthetic assembly line.

In vitro reconstitution of key biosynthetic transformations using simplified substrate analogues, combined with gene deletion and complementation experiments, enabled us to elucidate the origin of all structural differences in the C21 side chains of gladiolin and etnangien. The more saturated gladiolin side chain arises from a cis-acting enoylreductase (ER) domain in module 1 and in trans recruitment of a standalone ER to module 5 of the PKS. Remarkably, module 5 of the gladiolin PKS is intrinsically iterative in the absence of the standalone ER, accounting for the longer side chain in etnangien. These findings have important implications for biosynthetic engineering approaches to the creation of novel polyketide skeletons.

## Introduction

Polyketides are a large family of natural products that possess both impressive structural diversity and a broad range of biological activities. Several have found important applications as pharmaceuticals and agrochemicals, including erythromycin A (antibacterial), amphotericin B (antifungal), doxorubicin (anticancer agent), rapamycin (immunosuppressant), and avermectin (antiparasitic / insecticide).^1^

Etnangien **1** is a polyketide antibiotic produced by the myxobacterium *Sorangium cellulosum* that displays potent activity against a broad range of Gram-positive bacteria, including *Mycobacterium smegmatis* (Figure 1).^2^ However, it is highly unstable due to a light and acid-sensitive hexaene moiety in the C21 side chain. Gladiolin **2** is a structurally related antibiotic isolated from *Burkholderia gladioli* (Figure 1).^3^ Due to its shorter and more saturated C21 side chain, it has greater acid and light stability than etnangien.^4^ Gladiolin **2** has been reported to exhibit promising activity against drug-resistant clinical isolates of *Mycobacterium tuberlosis*.^4^

**Figure 1.**
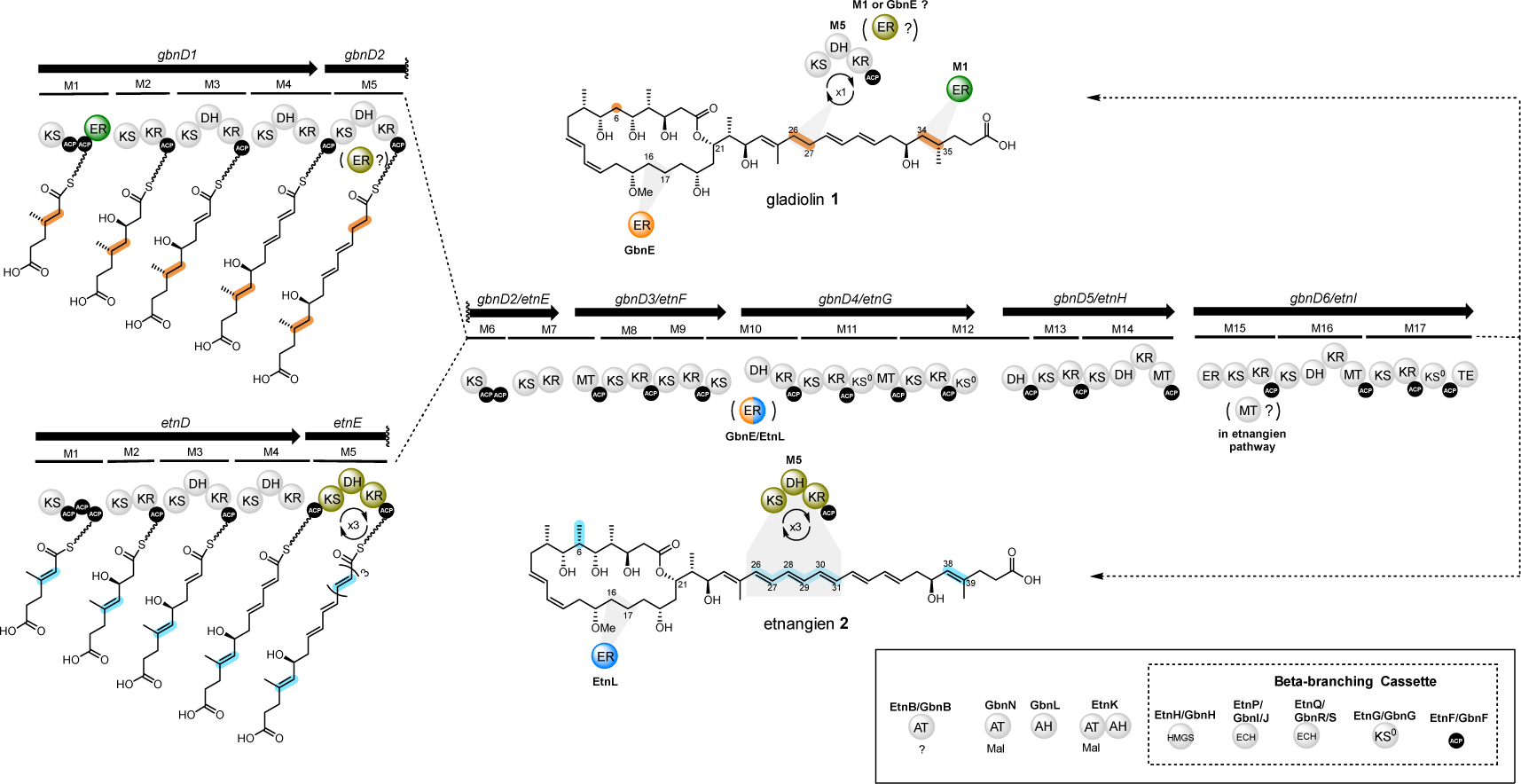
Comparison of the structures of gladiolin and etnangien and the architectures of the type I modular PKSs responsible for their assembly. Structural differences between gladiolin and etnangien are highlighted in pink and cyan, respectively. The architectures of the gladiolin and etnangien *trans*-AT PKSs are shown (modules are abbreviated M1, M2 etc.) with predicted intermediates in chain assembly attached to the ACP domains in each of the first five modules. The ER domain in module 1 of the gladiolin PKS, which replaces one of the ACP domains in module 1 of the etnangien PKS and is the sole architectural difference between the two assembly lines, is highlighted in green. The putatively iterative module 5 in the etnangien PKS and the additional ER domain required by module 5 of the gladiolin PKS are highlighted in gold. The *trans*-acting ERs GbnE and EtnL are highlighted in orange and blue, respectively. These are predicted to reduce the enoyl thioester intermediates attached to the ACP domains in module 10 of the respective PKS. Either the ER domain in module 1, or the *trans*-acting ER GbnE could be responsible for reduction of the enoyl thioester intermediate attached to module 5 of the gladiolin PKS. Other *trans*-acting enzymes involved in the assembly of gladiolin and etnangien are shown in the box (bottom right).

Most structurally complex polyketides are assembled in bacteria by type I modular polyketide synthases (PKSs). These are large multifunctional enzyme complexes consisting of a series of modules, each of which typically catalyzes one round of chain elongation and α / β-carbon modification.^5^ Each chain elongation cycle requires three domains: an acyl carrier protein (ACP), an acyltransferase (AT), and a ketosynthase (KS). Prior to initiation of chain assembly, ACP domains are post-translationally modified via attachment of a phosphopantetheine (Ppant) prosthetic group derived from coenzyme A. This is catalysed by a 4’-phosphopantethienyl transferase (PPtase) and results in conversion of the ACP domains from the *apo* to *holo* form. *Holo*-ACP domains use the Ppant ‘arm’ to shuttle covalently-bound acyl thioester intermediates between catalytic domains. Intermediates in chain assembly are translocated from the ACP domain in one module to the KS domain in the next via transthioesterification onto a conserved active site Cys residue. Decarboxylative condensation of the resulting thioester with an (alkyl)malonyl extender unit loaded onto the downstream ACP domain by an AT domain yields a β-keto thioester. Optional α/β-carbon modifying domains, including ketoreductase (KR), dehydratase (DH), enoylreductase (ER) and C/O-methyltransferase (MT) domains, further modify this intermediate.

Two architecturally and evolutionally distinct subclasses of type I modular PKSs have been identified: *cis*-AT and *trans*-AT. While *cis*-AT PKSs have AT domains that incorporate diverse extender units, including malonyl, methylmalonyl and ethylmalonyl thioesters, integrated into each module, *trans*-AT PKSs typically employ a single malonyl-CoA-specific standalone AT domain to supply extender units to each module.^6^ In addition to AT domains, *trans*-AT PKSs often recruit other *trans*-acting enzymes to modify the growing polyketide chain. For example, *trans*-acting ERs have been identified in several systems,^7,8,9^ including the kalimanticin PKS where BatK has been shown to reduce enoyl thioester intermediates in multiple modules.^10^ Other characteristic architectural features, including unusual domain orders, split modules and non-elongating modules have also been identified in *trans*-AT PKSs.^6^

While in bacteria type I modular PKSs are responsible for the assembly of most structurally complex polyketides, type I iterative PKSs are employed in fungi to assemble such structures.^11^ However, type I iterative PKSs are also found in bacteria^12^ and a recent study suggests they are more broadly distributed in bacterial species than originally assumed.^13^

Although type I PKSs generally function as either iterative or modular assembly lines, several examples of iterative module use in type I modular systems have been reported. These fall into two different categories: (1) an aberrant biosynthetic event, termed ‘stuttering’, yielding trace amounts of a by-product, such as ring expanded 6-deoxyerythonolide B^14^ and epothilones^15^, or (2) a programmed process yielding the primary product, such as aureothin^16^ (and structurally related products: neoaureothin^17^ and lutereotuclin^18^), stigmatellin,^19^ borellidin^20^ and azalomycin F.^21^ Programmed module iteration has inspired a rational engineering approach for structural diversification of polyketides: extending or reducing the chain length by switching on/off or reprograming module iteration. Early attempts to engineer a noniterative module from the erythromycin *cis*-AT PKSs to perform two rounds of chain elongation *in vitro* have been reported.^22^ To date only a few examples of programmed module iteration have been proposed in *trans*-AT PKSs, based on correlation of PKS architecture with the structure of the product.^6^ These include etnangien,^23^ 9-methylstreptimidone,^24^ lankacidin A,^25^ patellazole C,^26^ and nocardiosis associated polyketide (NOCAP)^27,28^ (Figure S1). Currently, little is known about the mechanisms underlying programmed module iteration in *trans*-AT PKS.

Gladiolin and etnangien (Figure 1) are a dramatic example of how deceptively similar modular PKS architectures can assemble structurally divergent products. These metabolites are assembled by *trans*-AT PKSs that differ by only a single domain – one of the ACP domains in module 1 of the etnangien PKS is replaced by an ER domain in the gladiolin PKS (Figure 2).^23,4^ Biosynthetic logic suggests most of the structural differences between the two compounds can be attributed to functional divergence in modules 1 and 5 of the PKSs^4, 23, 29^: (i) the saturated C34-C35 in gladiolin is posited to be a consequence of the ER domain in module 1 of the gladiolin PKS; and (ii) while the C26-C31 triene in etnangien is likely installed via three rounds of chain elongation, ketoreduction and dehydration by module 5 of the etnangien PKS, the saturated C26-C27 in gladiolin is proposed to result from a single round of chain elongation, ketoreduction, dehydration and enoylreduction by module 5 of the gladiolin PKS. Because the gladiolin and etnangien PKSs are highly homologous^4^, we postulated that module 5 of the etnangien and gladiolin PKSs may be intrinsically iterative, but that enoyl reduction after the first catalytic cycle prevents iteration in the latter. Intriguingly, however, module 5 of the gladiolin PKS does not contain an ER domain. We formulated two hypotheses to account for the cryptic enoylreduction in module 5 of the gladiolin PKS: (i) the ER domain in module 1 is bifunctional and catalyzes *inter*-modular reduction of the enoyl thioester intermediate generated by module 5, in addition to *intra*-modular reduction of the corresponding intermediate in module 1; or (ii) the putative *trans*-acting ER encoded by *gbnE* reduces enoyl thioester intermediates in modules 5 and 10 of the gladiolin PKS, whereas the corresponding ER encoded by *etnL* is only able to reduce an enoyl thioester attached to module 10 of the etnangien PKS.^4,29^

**Figure 2.**
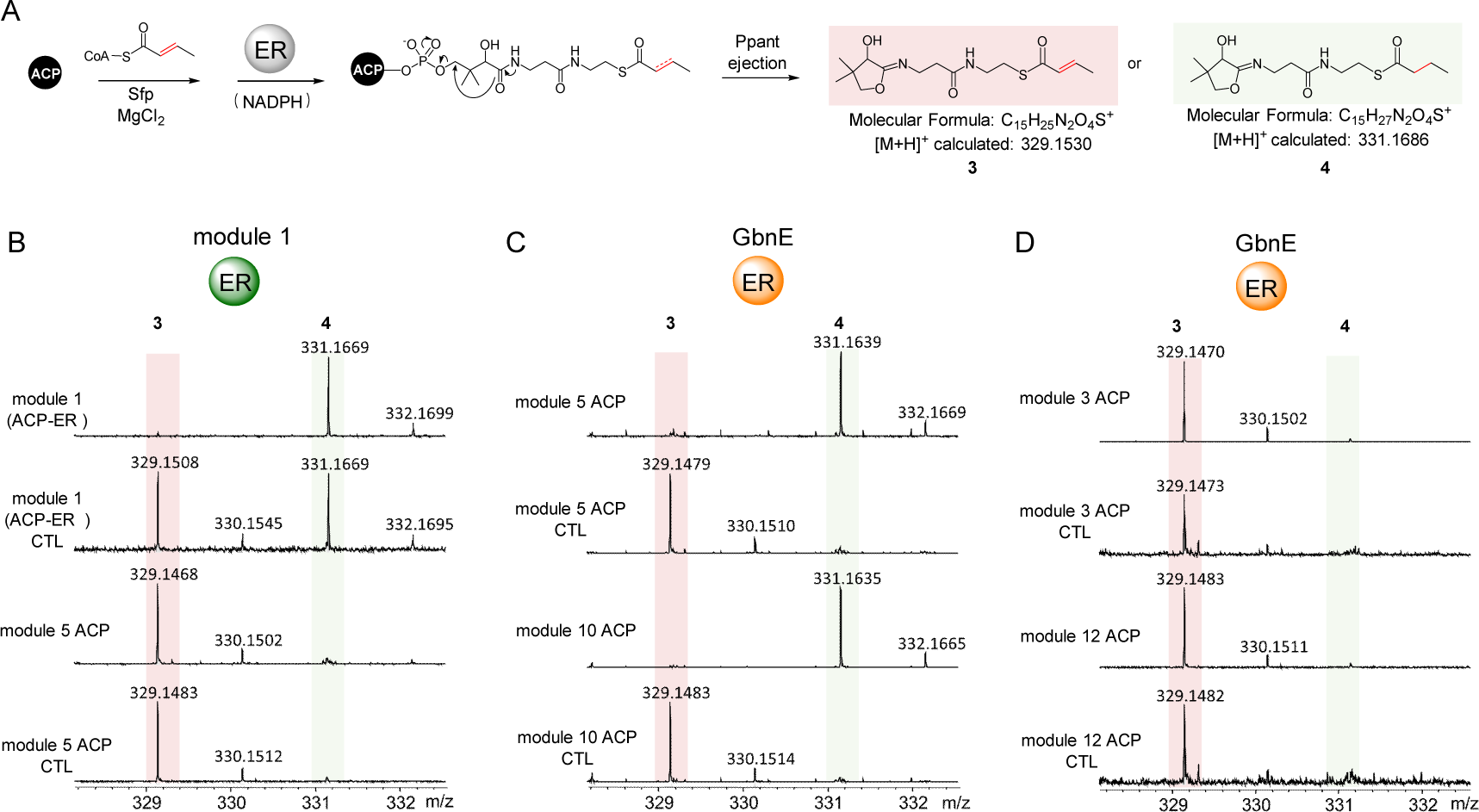
*In vitro* characterization of the activity of the module 1 ER domain and the *trans*-acting ER GbnE with a simplified mimic of enoyl thioester intermediates bound to various ACP domains of the gladiolin PKS. (A) Enoyl reduction assays used for characterization of the module 1 ER domain and GbnE. Negative control reactions omitted the module 1 ER domain or GbnE, except when the crotonyl-*holo*-module 1 ACP-ER didomain was employed, where NADPH was omitted. Structures and calculated *m/z* values for the Ppant ejection ions arising from the substrate and product of enoyl reduction are shown. (B) Ppant ejection ions observed from intact MS analysis of crotonylated module 1 *holo*-ACP-ER didomain following incubation with NADPH and the crotonylated *holo*-ACP domain excised from module 5 following incubation with the excised module 1 ER domain and NADPH. (C) Ppant ejection ions observed from intact MS analysis of the crotonylated *holo*-ACP domains excised from modules 5 and 10 following incubation with GbnE and NADPH. (D) Ppant ejection ions observed from intact MS analysis of crotonylated *holo*-ACP domains excised from modules 3 and 12 following incubation with GbnE and NADPH.

Here we report an extensive set of biochemical and genetic experiments that reveal the molecular basis for assembly of the C26-C38 region of gladiolin, which is significantly shorter and more saturated than the corresponding C26-C42 region of etnangien. First, we show that the ER domain in module 1 of the gladiolin PKS is responsible for saturation of C34-C35. Second, we demonstrate that GbnE and EtnL are *trans*-acting ERs able to reduce ACP-bound enoyl thioester intermediates in both modules 5 and 10 of the gladiolin PKS, but only module 10 of the etnangien PKS. Third, we show module 5 of the gladiolin PKS is intrinsically iterative, but that GbnE-catalyzed enoylreduction after one round of chain elongation, ketoreduction, and dehydration suppresses iteration. Fourth, we show the KS domain in module 6 of the gladiolin PKS preferentially accepts saturated thioesters over α, β-unsaturated counterparts, promoting module 5 iteration in the absence of enoyl reduction. Together these data provide significant new insight into how Nature reprograms complex metabolite assembly by *trans*-AT PKSs.

## Results and discussion

### GbnE is the cryptic catalyst for enoyl reduction in module 5 of the gladiolin PKS

To investigate whether the ER domain in module 1 of the gladiolin PKS, or the putative *trans*-acting ER encoded by *gbnE* is responsible for reduction of the enoyl thioester intermediate attached to the ACP domain in module 5, we investigated the catalytic activity of the excised module 1 ER domain and GbnE *in vitro*. The module 1 ACP-ER didomain, the module 1 ER domain, the module 5 ACP domain and the module 10 ACP domain were overproduced as soluble *N*-terminal His6 fusion proteins in *E. coli*. Due to problems with obtaining sufficient quantities of soluble GbnE using this approach, we employed an N-terminal His6-SUMO (small ubiquitin-related modifier) fusion. The resulting recombinant proteins were purified using immobilized metal-ion affinity chromatography (IMAC). An additional size exclusion chromatography step was used to further purify GbnE. The identity of each purified protein was confirmed by SDS-PAGE and ESI-Q-TOF-MS analyses (Figure S2). The latter confirmed that the module 5 and 10 ACP domains and the module 1 ACP-ER didomain were all in the *apo* form. GbnE exhibited a notable yellow colour, indicative of a bound flavin cofactor. UHPLC-Q-TOF-MS comparison of an extract from denatured GbnE with an authentic standard confirmed this is flavin mononucleotide (FMN) (Figure S3).

A crotonyl thioester was used as a simplified mimic of the enoyl thioester intermediates attached to the ACP domains in modules 1, 5 and 10 during gladiolin assembly. The *apo*-ACP domains were converted to the crotonylated *holo*-forms by incubating with crotonyl-coenzyme A and the substrate tolerant phosphopantetheinyl transferase Sfp (Figure 2A).^30^ A 409 Da mass increase relative to the corresponding *apo* protein was observed in intact UHPLC-ESI-Q-TOF-MS analyses, confirming that crotonyl-Ppant had been loaded onto each ACP domain (Figure S4).

To investigate the function of the module 1 ER domain, the crotonylated module 1 *holo*-ACP-ER didomain was incubated with NADPH, and the crotonylated module 5 *holo*-ACP domain was incubated with a mixture of the module 1 ER domain and NADPH. Because the molecular weights of these proteins are too large to distinguish a 2 Da mass shift by intact UHPLC-ESI-Q-TOF-MS, Ppant ejection was used to determine the outcome of these reactions (Figure 2A).^31^ Only the crotonyl units attached to the module 1 ACP-ER didomain and the module 1 ACP domain were reduced (Figures 2B). No reduction of the crotonylated module 5 *holo*-ACP domain was observed (Figure 2B). Approximately 50% reduction of the crotonyl group attached to the module 1 ACP-ER di-domain was observed even when NADPH was omitted. This is likely because significant quantities of unreacted NADPH remain bound to the active site of the ER domain in the ACP-ER didomain construct throughout purification. These results show that the module 1 ER domain can catalyze reduction of enoyl thioesters attached to the second ACP domain in module 1 of the gladiolin PKS, but not the ACP domain from module 5.

To examine the role played by GbnE, the crotonylated *holo*-ACP domains from modules 5 and 10 were incubated with GbnE and NADPH. The Ppant ejection ions observed in UHPLC-ESI-Q-TOF-MS/MS analyses of the reaction mixtures showed the crotonyl group underwent reduction in both cases (Figure 2C). This indicates that GbnE catalyzes in-*trans* reduction of the enoyl thioester intermediates generated by modules 5 and 10 during gladiolin assembly. To probe whether the ACP domains are the principal specificity determinant for recruitment of GbnE to these modules, we performed analogous assays with excised ACP domains from modules 3 and 12 of the PKS. Module 3 produces an enoyl thioester intermediate that is predicted not to undergo further reduction, because a C-C double bond is found in the corresponding C30-C31 region of gladiolin. Module 12 produces a β-hydroxy thioester that undergoes a further round of chain elongation and ketoreduction on module 13. The resulting β, δ-dihydroxy thioester is converted to the corresponding *E*, *Z*-diene by the DH-like domain in module 13.^32^ The Ppant ejection ions observed in UHPLC-ESI-Q-TOF-MS/MS analyses of these reaction mixtures confirmed that a crotonyl thioester attached to these ACP domains cannot be reduced by GbnE (Figure 2D).

Taken together, these data are consistent with in-*cis* reduction of the enoyl thioester intermediate attached to the module 1 ACP domain by the adjacent ER domain and in-*trans* reduction of the enoyl thioester intermediates attached to the module 5 and 10 ACP domains by GbnE.

### GbnE plays a key role in gladiolin biosynthesis *in vivo*

To further substantiate the proposed role of GbnE in gladiolin biosynthesis, we constructed an in-frame deletion in *gbnE* in the gladiolin producer *B. gladioli* BCC1622 (Figure S5).^33^ UHPLC-ESI-HRMS analysis of extracts from the mutant showed the production of gladiolin was significantly diminished but not completely abolished (Figure 3A). Genetic complementation of the *gbnE* mutant using a derivative of pMLBAD with *gbnE* under the control of an arabinose-inducible promotor (Figure S5) restored gladiolin production to near-wild type levels (Figure 3A). These results confirmed that GbnE plays an important role in gladiolin biosynthesis, but suggested its function could be partially complemented by another enzyme *in vivo*. The *gdsB* gene in the gladiostatin biosynthetic gene cluster in *B. gladioli* BCC1622 encodes a protein predicted to contain acyl hydrolase (AH), AT, and ER domains.^34^ The ER domain of GdsB shares 57% sequence identity with GbnE. A double knockout mutant of *B. gladioli* BCC1622, containing in frame deletions in both *gbnE* and the ER domain of *gdsB* was therefore created (Figure S5). Gladiolin production was completely abolished in this mutant (Figure 3A), confirming that the GdsB ER domain can partially compensate for the loss of in-*trans* ER activity in the *gbnE* mutant.

**Figure 3.**
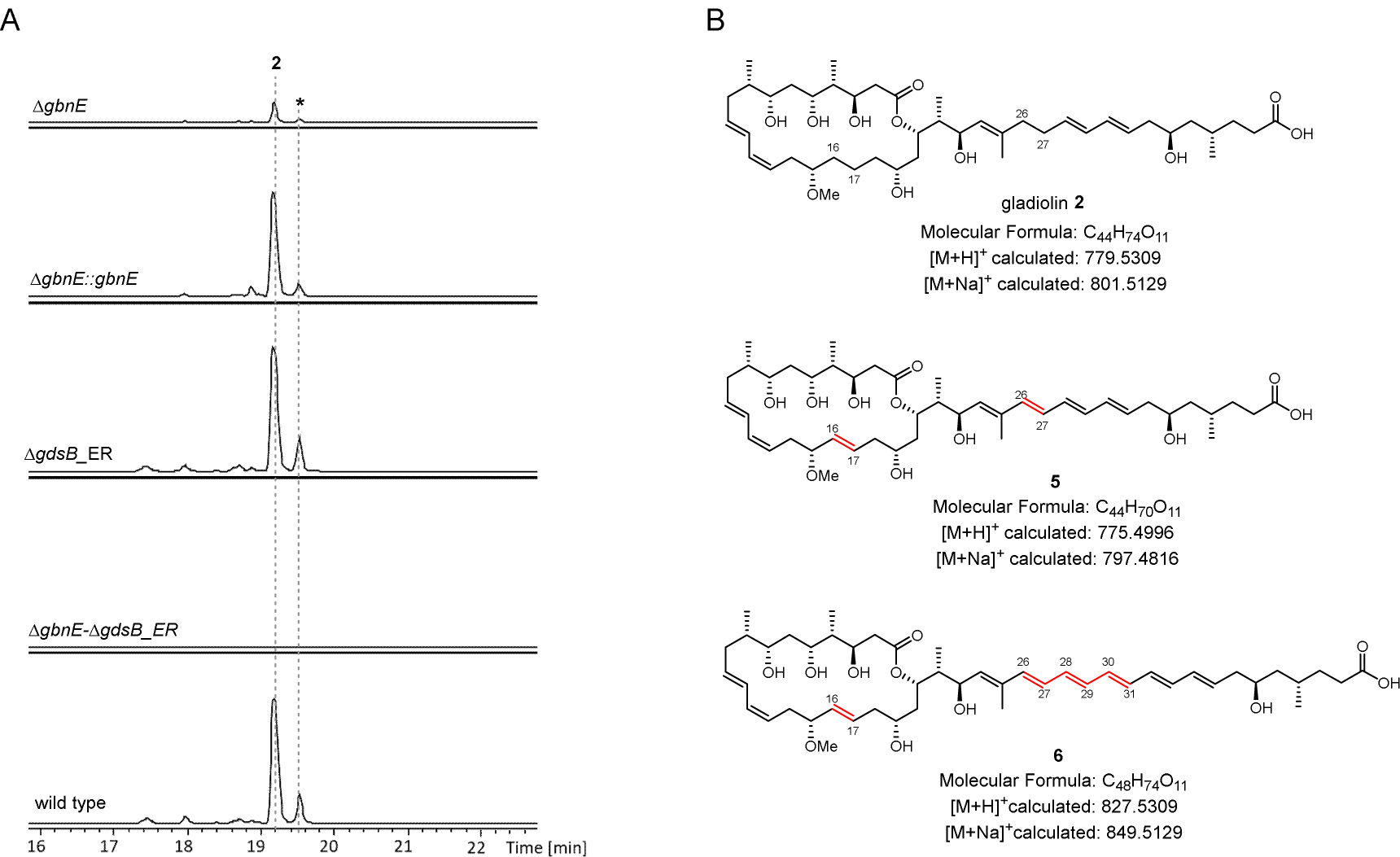
UHPLC-ESI-HRMS analysis of the production of gladiolin and derivatives and in *gbnE* and *gdsB*_ER in-frame deletion mutants and the complemented *gbnE* mutant. (A) Extract ion chromatograms corresponding to the [M+H]^+^ and [M+Na]^+^ ions for **2**, **5** and **6** from wild type *B. gladioli* BCC1622, the *gbnE* mutant, the *gdsB_*ER mutant, the *gbnE*-*gdsB_*ER double mutant and the *gbn*E mutant complemented by in *trans* expression of *gbnE* under the control of an arabinose-inducible promoter. The asterisk denotes *iso*-gladiolin, resulting from a spontaneous rearrangement of gladiolin,^3^ involving migration of the C1 acyl group to the C23 hydroxyl group. (B) Structural comparison of gladiolin (**2**) with the gladiolin derivatives **5** and **6** that could potentially be produced by the *gbnE* mutant and the *gbnE*-*gdsB_*ER double mutant.

No gladiolin-related shunt metabolites could be observed in either of the mutants including compounds with *m/z* values corresponding to the [M+H]^+^ ions for derivative **5**, containing C16-C17 and C26-C27 double bonds (expected to result from abrogation of ER activity in modules 5 and10), or derivative **6**, containing a C16-C17 double bond and a C26-C31 triene (expected to result from three iterations of module 5, in addition to abrogation of ER activity in modules 5 and10) (Figure 3B). Overall, these data show that a *trans*-acting ER is an essential component of the machinery for gladiolin assembly.

### Module 5 of the gladiolin PKS is intrinsically iterative

We next investigated whether module 5 of the gladiolin PKS, which contains KS, DH, KR and ACP domains, can catalyze multiple rounds of chain elongation, ketoreduction, and dehydration and whether enoyl reduction by GbnE can suppress this. This module was overproduced in *E. coli* as an N-terminal His6 fusion and purified by Ni-NTA affinity and size exclusion chromatography. UHPLC-ESI-Q-TOF-MS analysis showed the ACP domain of this construct was in the unmodified *apo* form (Figure S2). Unfortunately, attempts to overproduce GbnN, the putative *trans*-acting AT that loads malonyl extender units onto the ACP domains of the gladiolin PKS, in soluble form were unsuccessful. We therefore overproduced and purified the C-terminal domain of EtnK, the analogous AT from the etnangien PKS (65% sequence similarity; Figure S2).

N-acetylcysteamine (NAC) thioesters can serve as effective mimics of ACP-bound intermediates in polyketide chain assembly, enabling acylation of the active site Cys residue of cognate KS domains *in vivo* and *in vitro*.^35,36^ We thus employed 2,4-hexadienoyl NAC thioester **7** as a structurally simplified surrogate of the intermediate bound to module 4 of the gladiolin PKS. Purified recombinant module 5 was first incubated with NAC thioester **7** to prime the KS active site Cys residue with the 2, 4-hexadienoyl group, then malonyl-CoA / Sfp (to load the extender unit onto the ACP domain) and NADPH (the co-substrate for the KR domain) were added (Figure 4A). These reagents are sufficient for one round of chain elongation, ketoreduction and dehydration. To enable multiple iterations of this process, EtnK was also added to recharge the *holo*-ACP domain with a malonyl extender unit after the product from the first iteration has been transferred back onto the KS domain. After 20h incubation, this reaction mixture was yellow, indicating a polyene had been formed, whereas a control reaction containing a C209A mutant of module 5, in which the active site Cys of the KS domain was replaced by Ala, was colorless.

**Figure 4.**
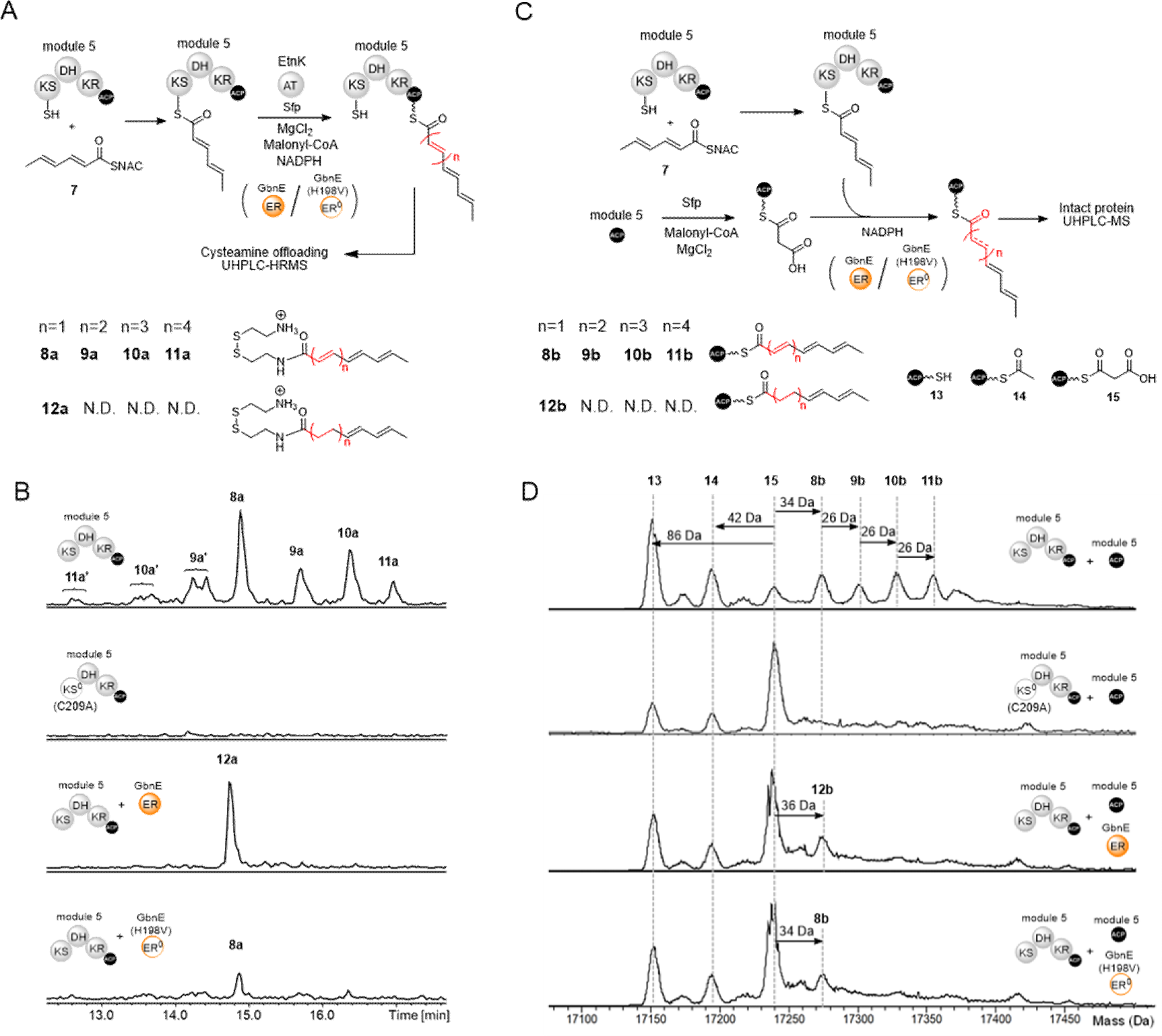
*In vitro* reconstitution of module 5 from the gladiolin PKS in the presence and absence of GbnE and an H198V mutant. (A) Scheme illustrating the design of the module 5 reconstitution assays. The KS and ACP domains are primed using 2, 4-hexadienoyl thioester **7** and malonyl-CoA / Sfp, respectively. The *trans*-acting AT domain from the EtnK subunit of the etnangien polyketide synthase is included to enable reloading of malonyl-CoA onto the *holo*-ACP domain after the first round of chain elongation, ketoreduction, dehydration and back transfer onto the active Cys residue of the KS domain. The effect of the *trans*-acting ER GbnE on iteration was examined by comparing products formed in the presence / absence of GnbE and the presence of a catalytically inactive H198V mutant of GbnE. The products formed were offloaded using cysteamine. (B) Extracted ion chromatograms at *m/z* values corresponding to the [M+H]^+^ ions for cysteamine offloading adducts **8a**, **9a**, **10a**, **11a** and **12a** from UHPLC-ESI-Q-TOF-MS analyses of reactions in the presence and absence of GbnE, and the presence of the H198V mutant of GbnE. No products are observed in a control reaction lacking GbnE and employing a C209A mutant of module 5, which has a catalytically inactive KS domain. **9a′**, **10a′** and **11a′** are hypothesized to be stereoisomers of **9a, 10a** and **11a** resulting from *E* to *Z* configurational isomerization of one or more double bonds in the polyene. (C) Scheme illustrating the design of module 5 plus module 5 ACP domain reconstitution assays. The assays follow similar procedures to the module 5 reconstitution reactions except that EtnK is replaced with the excised module 5 ACP domain loaded with a malonyl unit. The products attached to the excised module 5 ACP domain were identified by UHPLC-ESI-Q-TOF-MS analyses of the intact protein. (D) Deconvoluted mass spectra of the excised module 5 ACP domain following *in vitro* reconstitution with module 5 (top), module 5(C209A) as a negative control (2nd from top), module 5 + GbnE (3rd from top) and module 5 + GbnE(H198V) (bottom).

We initially attempted to monitor product formation using intact protein mass spectrometry. However, due to the large size of module 5 (∼160 kDa), there was insufficient resolution in the deconvoluted mass spectra to distinguish the mass increases resulting from each round of chain elongation and subsequent modification. We thus used cysteamine to offload the products of the reaction and UHPLC-ESI-Q-TOF-MS to characterize the resulting adducts.^37^ This led to the identification of species with molecular formulae corresponding to 1,3,5-hexatrienoyl, 1,3,5,7-octatetraenoyl, 1,3,5,7,9-decapentaenoyl and 1,3,5,7,9,11-dodecahexaenoyl adducts **8a-11a** (Figure 4B and Figure S7). None of these adducts were observed in the control reaction.

To confirm that these products result from module iteration, the assay was repeated using the module 5 *apo*-protein and a two-fold excess of the excised malonylated *holo*-ACP domain from module 5, in the absence of EtnK and exogenous malonyl-CoA (Figure 4C). A similar approach has previously been used to interrogate the mechanism of combinatorial chain elongation by the quartromicin polyketide synthase, where it was shown that an excised ACP domain can interact productively in *trans* with the KS and AT domains of the *apo*-QmnA3 protein.^21,38^ The small size of the excised ACP domain compared with the intact module enables direct observation of the intermediates accumulated using intact protein MS. Deconvoluted mass spectra from UHPLC-ESI-Q-TOF-MS analysis of the excised module 5 ACP domain revealed adducts **8b-11b** were gradually formed, with masses corresponding to 1-4 rounds of chain elongation, ketoreduction and dehydration, ultimately resulting in the 1,3,5,7,9,11-dodecahexaenoyl thioester **11b** (Figures 4D and S7). The acetylated excised *holo*-ACP domain **14** was also observed, presumably formed by either spontaneous or KS domain-catalyzed decarboxylation of the malonyl thioester. The ACP-bound adducts produced in this assay were offloaded with cysteamine, and UHPLC-ESI-Q-TOF-MS analyses were used to confirm their identity (Figure S8) In a control assay, using a C209A mutant of the *apo*-module 5, only the acetylated species **14** was formed (Figure 4D). Taken together, these observations demonstrate that module 5 of the gladiolin PKS is intrinsically iterative.

### GbnE suppresses iteration of module 5

To investigate whether GbnE can suppress the intrinsically iterative nature of module 5, we repeated the assays involving cysteamine offloading of ACP-bound intermediates from the intact module and MS monitoring of intermediates accumulated on the excised ACP domain in the presence of GbnE. In both cases, the reaction mixtures were colorless and only a single round of chain elongation, ketoreduction, dehydration and enoyl reduction was observed (Figures 4B, 4C and S8). No products resulting from further rounds of chain elongation and subsequent modification were detected. These data show that GbnE suppresses iteration of module 5. This could be because interactions between GbnE and module 5, prevent the next round of chain elongation and modification, or because the α, β-saturated thioester product of GbnE-catalyzed enoyl reduction is a poor substrate for transacylation back onto the module 5 KS domain compared to the α, β-unsaturated thioester intermediate.

To investigate the effect of interactions between GbnE and module 5 on module iteration, we sought to generate a catalytically inactive, but correctly folded, GbnE mutant. Sequence alignment of *trans*-acting ERs with the structurally and phylogenetically related homologue FabK, the FMN-dependent enoyl-ACP reductase involved in bacterial fatty acid biosynthesis, revealed the catalytically important His residue (H144) in the GG<u>H</u>T/I motif of FabK is highly conserved in *trans*-acting ERs (Figure S9).^39^ We therefore mutated it to Val in GbnE, generating GbnE-H198V. Purified GbnE-H198V was yellow, indicating FMN is bound and the protein is correctly folded (Figure S2). No reduction was observed following incubation of GbnE-H198V with the 2,4-hexadienoylated module 5 *holo*-ACP domain, confirming the mutation abolishes enoylreductase activity (Figure S10). Remarkably, when GbnE was replaced with GbnE-H198V in the module iteration assays, only a single round of chain elongation, ketoreduction and dehydration was observed, both when using cysteamine off-loading and the excised module 5 ACP domain in *trans* to monitor product formation (Figures 4B and 4C). This demonstrated that interaction between GbnE and module 5 is the minimum epitope for suppression of module iteration – reduction of the ACP-bound enoyl thioester intermediate is not required.

Interestingly, we noted that the amount of GbnE required to suppress module iteration was surprisingly low (Figure S7). Even when module 5 was in a ten-fold excess relative to GbnE, no iteration was observed. When we increased the amount of GbnE, a product resulting from one round of chain elongation with no ketoreduction or dehydration **16** was observed. At a ratio of module 5: excised module 5 ACP domain: GbnE of 1:2:0.5 this became the major product of the reaction (Figure S7).

### EtnL can substitute for GbnE in gladiolin biosynthesis

Having established that GbnE suppresses the iteration of module 5 in the gladiolin PKS, we next examined whether EtnL, the equivalent *trans*-acting ER encoded by the etnangien BGC, behaves similarly. To interrogate the proposed enoylreductase function of EtnL in etnangien biosynthesis, we overproduced and purified EtnL as an N-terminal MBP fusion protein and the excised ACP domains from modules 5 and 10 of the etnangien PKS as N-terminal His_6_ fusion proteins. Enoyl reduction assays performed as described for GbnE showed that EtnL can only reduce enoyl thioesters attached to the module 10 ACP domain of the etnangien PKS, and not the module 5 ACP domain (Figure 5A). In contract, EtnL reduced enoyl thioesters attached to the excised ACP domains from both module 5 and module 10 of gladiolin PKS (Figure 5A). Similarly, substitution of GbnE with EtnL in the gladiolin module 5 iteration assays yielded only products resulting from a single round of chain elongation, ketoreduction, dehydration and enoyl reduction (Figure 5C and 5D); no products of iterative chain elongation were observed.

**Figure 5.**
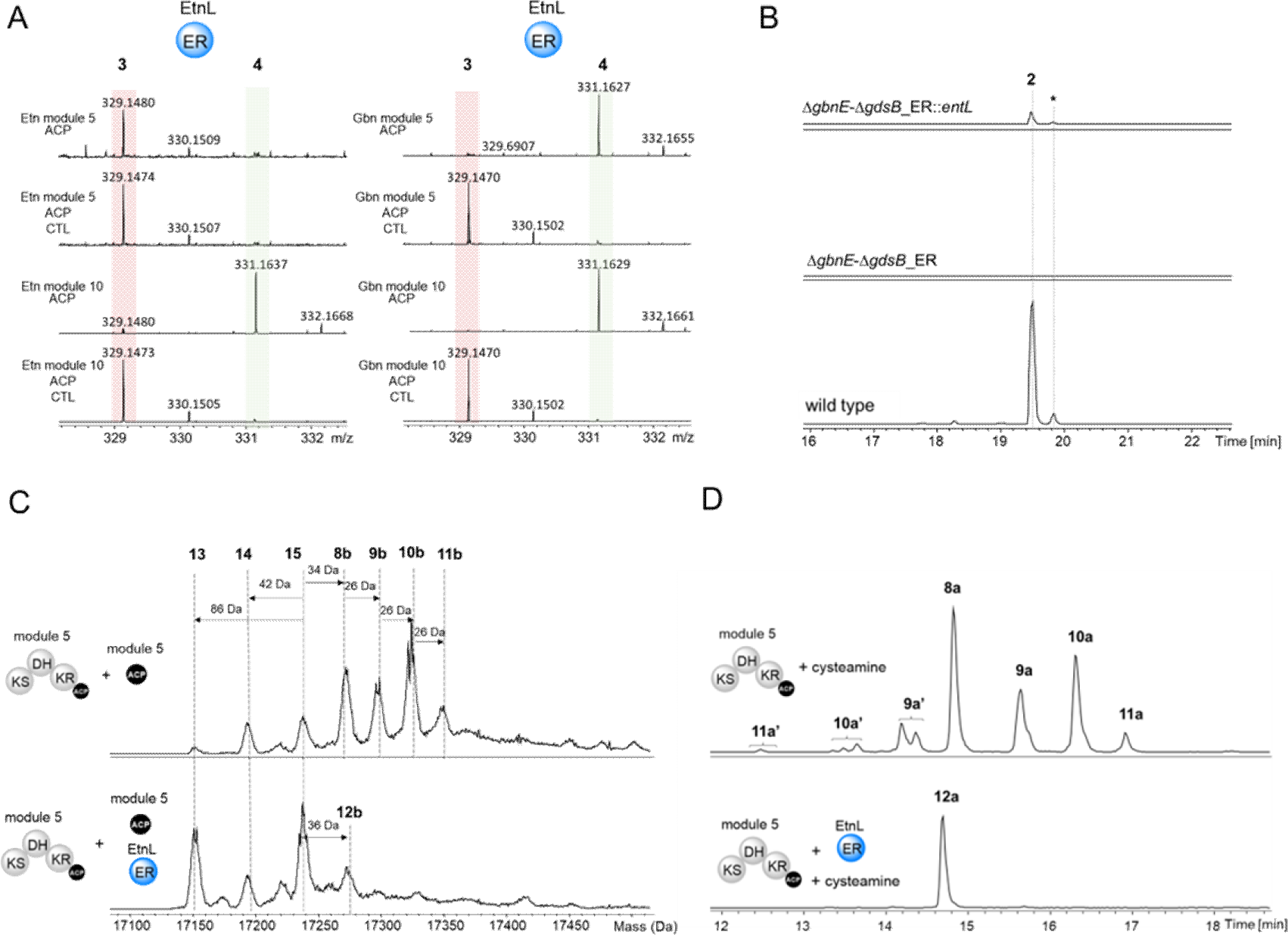
Characterization of EtnL function *in vitro* and *in vivo*. (A) Ppant ejection ions observed in UHPLC-ESI-Q-ToF-MS/MS analyses of the crotonylated *holo*-ACP domains excised from modules 5 and 10 of the etnangien PKS (left) and modules 5 and 10 of the gladiolin PKS (right), following incubation with EtnL and NADPH. (B) Extracted ion chromatograms at *m/z* values corresponding to [M+H]^+^ for gladiolin **2** and gladiolin derivatives **3** and **4** from UHPLC-ESI-Q-TOF-MS analyses of extracts from *B. gladioli* BCC1622 (wild type), the *gbnE*-*gdsB_ER* double mutant, and the *gbnE*-*gdsB_ER* double mutant complemented by in *trans* expression of *etnL* under the control of an arabinose inducible promoter. The asterisk denotes *iso*-gladiolin, resulting from a spontaneous rearrangement of gladiolin,^3^ involving migration of the C1 acyl group to the C23 hydroxyl group. (C) Deconvoluted mass spectra from UHPLC-ESI-Q-ToF-MS analyses of the excised ACP domain from module 5 of the gladiolin PKS, following incubation with intact module 5 from the gladiolin PKS, 2, 4-hexadienoyl thioester **7**, malonyl-CoA, Sfp and EtnK in the absence (top) and presence (bottom) of EtnL. (D) Extracted ion chromatograms at *m/z* values corresponding to [M+H]^+^ ions for cysteamine offloading adducts **8a**, **9a**, **10a**, **11a** and **12a** from UHPLC-ESI-Q-ToF-MS analyses of incubation mixtures containing module 5 from the gladiolin PKS, 2, 4-hexadienoyl thioester **7**, malonyl-CoA, Sfp and EtnK in the absence (top) and presence (bottom) of EtnL following treatment with cysteamine. **9a′**, **10a′** and **11a′** are hypothesized to be the stereoisomers of **9a, 10a** and **11a**, respectively, resulting from *E* to *Z* configurational isomerization of one or more double bond in the polyene.

Overall, these results show that EtnL can play a functionally equivalent role to GbnE in gladiolin biosynthesis. As further proof of this, gladiolin production was restored and no shunt metabolites were observed when *etnL* was used to complement the *B. gladioli* BCC1622_Δ*gbnE*Δ*gdsB*_ER mutant, albeit to only a modest level, which could be due to sub-optimal expression of *etnL* (from *Sorangium*) in *B. gladioli* (Figure 5B).

### The module 6 ketosynthase domain prefers α, β-saturated thioester substrates

Although the above data demonstrate that excised module 5 from the gladiolin PKS is intrinsically iterative and iteration is supressed by the *trans*-acting ER, in the intact assembly line the module 6 KS domain may also contribute to suppression of iteration. This KS domain is fused to the C-terminus of module 5 in the GbnD2 subunit and can accept intermediates bound to the module 5 ACP domain via transacylation onto its active site Cys residue. Thus, the module 5 and 6 KS domains may compete for intermediates bound to the module 5 ACP domain and whether back or forward transfer of intermediates is favoured could be influenced by the substrate preference of each KS domain in the transacylation step. The above experiments indicate that the module 5 KS domain greatly prefers α, β-unsaturated over α, β-saturated thioesters in the back transfer (or subsequent chain elongation) reaction. We thus sought to establish the substrate preference of the module 6 KS domain in both the transacylation and chain elongation steps.

*N*-terminal His_6_ fusions of both the KS-ACP didomain and the KS domain were found to be insoluble. Thus, we overproduced the entire excised module 6 (KS-ACP-ACP tri-domain) as an N-terminal His_6_-fusion and mutated the conserved Ser residue in the second ACP domain to Ala. This mutation ensures the protein can only bear a single phosphopantetheine prosthetic group, which simplifies intact protein mass spectrometry analyses.

We first probed the transacylation substrate specificity of the module 6 KS domain towards simplified α, β-unsaturated and α, β-saturated analogues of the native biosynthetic intermediates before and after GbnE-catalysed enoyl reduction. The S941A mutant of module 6 was incubated with the module 5 *apo-*ACP domain that had been loaded with (2*E*, 4*E*)-2,4-hexadienoyl-Ppant or (*E*)-4-hexenoyl-Ppant thioesters via incubation with the corresponding pantetheine thioesters (**17** and **18**), Sfp, CoaA, CoaC and CoaD (Figure 6A, Figure S11). The amount of chain transacylation onto the module 6 KS domain was monitored by comparing the ratio of module 5 S-acyl to *holo*-ACP domain in each case. The deconvoluted mass spectra of the ACP domains from each reaction showed that, after 8 hours incubation with the module 6 S941A mutant, significantly more of the 4-hexenoyl thioester had been transferred to the KS domain than the 2,4 hexadienoyl thioester (Figure 6B). Indeed, compared to a control reaction lacking the module 6 S914A mutant, less than 5% of the 2,4 hexadienoyl thioester had been transferred. This clearly demonstrates the preference of the module 6 KS domain for α, β-saturated over α, β-unsaturated substrates in the transacylation step. We next investigated the specificity of the chain elongation step by using the NAC thioesters of (2*E*, 4*E*)-2, 4-hexadienoic acid (**7**) and (*E*)-4-hexenoic acid (**19**) to acylate the active site Cys residue of the KS domain then loading malonyl-ppant onto the *apo*-ACP domain and using intact protein MS to monitor chain elongation. These experiments showed that the module 6 KS domain does not prefer the 4-hexenoyl thioester over the 2,4-hexadienoyl thioester in the chain elongation step (Figure S12).

**Figure 6.**
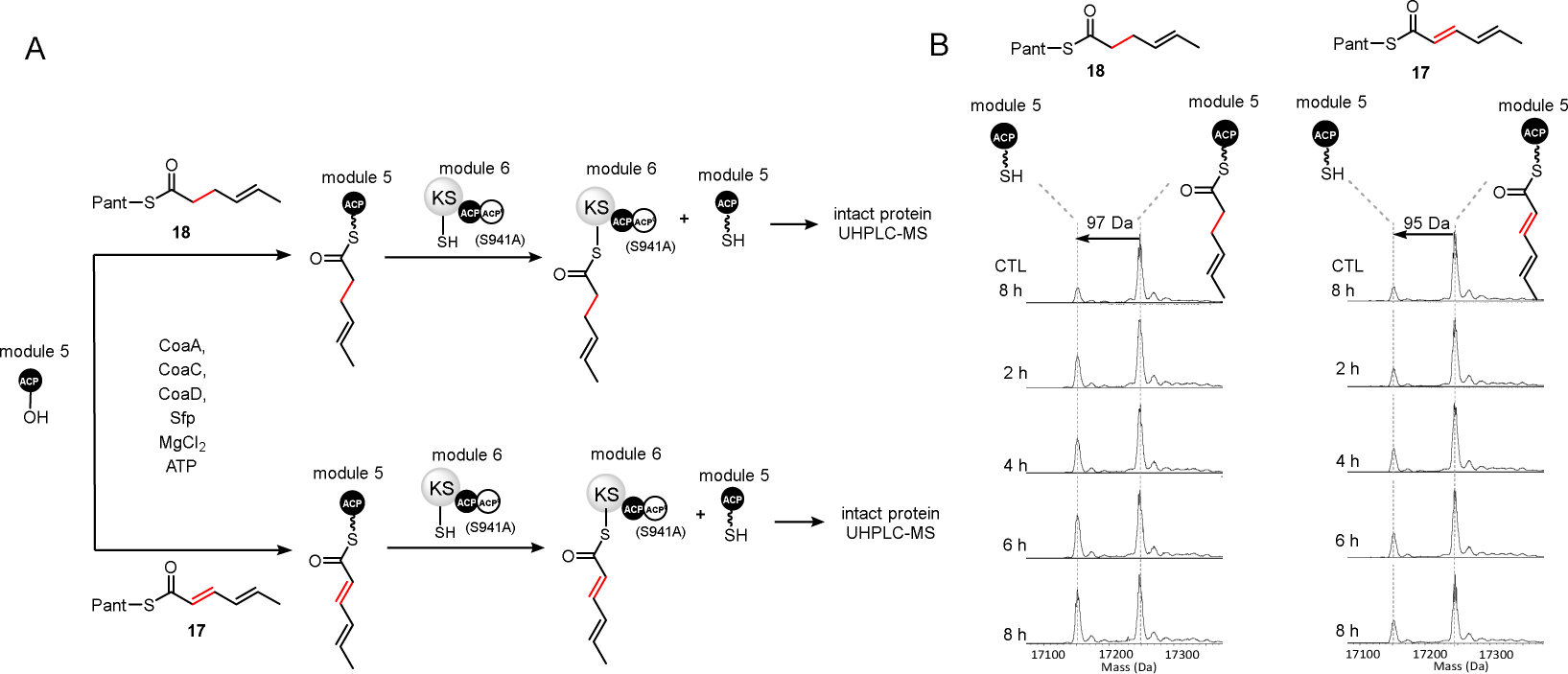
Preference of the module 6 KS domain towards simplified mimics of the α, β-saturated and α, β-unsaturated intermediates attached to the module 5 ACP domain during gladiolin chain assembly. (A) Scheme illustrating the design of the module 6 KS domain substrate preference assays. The pantetheine thioesters of (2*E*, 4*E*)-2, 4-hexadienoate and (4*E*)-4-hexenoate, **17** and **18** respectively, are converted to the corresponding coenzyme A thioesters and loaded onto the excised ACP domain from module 5 of the gladiolin PKS using purified recombinant CoaA, CoaC, CoaD and Sfp. The resulting ACP thioesters are incubated with an S941A mutant of purified recombinant module 6 from the gladiolin PKS and the extent of module 5 *holo*-ACP formation was determined at various time points using UHPLC-ESI-Q-ToF-MS analysis of the intact protein. (B) Deconvoluted mass spectra of the 4-hexaenoylated (left) and 2,4-hexadienoylated (right) module 5 *holo*-ACP domain following incubation with a S941A mutant of module 6 from the gladiolin PKS. Deacylation of the module 5 *holo*-ACP domain was used to monitor transfer onto the module 6 KS domain. A negative control reaction omitting module 6(S941A) was conducted in each case.

Taken together, our results indicate that the substrate specificity of the module 6 KS domain contributes to the overall programming of chain assembly by the gladiolin PKS by ensuring the α, β-unsaturated thioester intermediate attached to the module 5 ACP domain does not get translocated onto the downstream KS domain until it has undergone GbnE-catalysed enoyl reduction. They also explain why a lack of EtnL-catalysed enoylreduction in module 5 of the etnangien PKS causes this module to iterate, resulting in incorporation of the C26-C31 triene into etnangien.

### Conclusions

The data reported here indicate that the significant differences in length and degree of unsaturation between the gladiolin and etnangien C21 side chains result primarily from (i) substitution of an ACP domain in module 1 of the etnangien PKS by an ER domain in the gladiolin PKS, resulting in reduction of the β-methyl-α, β-unsaturated thioester intermediate attached to the module 1 ACP domains in gladiolin biosynthesis; (ii) the intrinsically iterative nature of module 5 in the gladiolin and etnangien PKSs, enabling three rounds of chain elongation, ketoreduction and dehydration, leading to formation of a 1, 3, 5-hexatrienoyl thioester intermediate by the latter; and (iii) differential recruitment of a *trans*-acting ER to modules 5 and 10 of the gladiolin PKS, but only module 10 of the etnangien PKS, which suppresses iteration of module 5 in the former.

Interestingly, the KS domain in module 6 of the gladiolin PKS prefers an α, β-saturated over an α, β-unsaturated acyl donor, suggesting it may act as a gatekeeper to ensure the enoyl thioester attached to the module 5 ACP domain is reduced prior to further chain elongation. It is reasonable to assume that the KS domain in module 6 of the etnangien PKS has a similar acyl donor preference. This would suppress transfer of the enoyl thioester intermediate initially formed by module 5 of the entangien PKS onto the module 6 KS domain, promoting back transfer onto the module 5 KS domain, leading to further rounds of chain elongation ketoreduction and dehydration. Once a third iteration of this cycle has been completed, the module 6 KS domain is presumably able to readily accept the resulting 1, 3, 5-hexatrienoyl thioester enabling assembly of the etnangien polyketide chain to be completed.

In addition to *trans*-AT PKSs, iteration is a feature of several other type I PKS systems, including bacterial *cis*-AT PKSs and both bacterial and fungal iterative PKSs.^40^ Among these, iterative fungal PKSs involved in the assembly of lovastatin, tennelin, and desmethylbassianin are relevant to the findings reported here, because *trans*-acting ERs have been shown to play a key role in programming of chain assembly.^41,42^ However, it is important to note that the *trans*-acting ERs associated with fungal iterative PKSs belong to a different family to their counterparts in *trans*-AT PKSs.^42^ Intriguingly, our data show that catalytically inactive GbnE is still able to suppress iteration of module 5 in the gladiolin PKS, indicating that association of the ER with the module, rather reduction of the α, β-unsaturated thioester intermediate, controls iterative module use in *trans*-AT PKSs. Further investigations will be required to fully elucidate the nature of the interaction between GbnE and module 5 of the gladiolin PKS. Our data clearly indicate that the nature of the ACP domain is a key determinant of *trans*-acting ER recruitment to *trans*-AT PKS modules, but it is unclear whether interactions with other domains also play a role. A more complete understanding of how *trans*-acting ERs are recruited and suppress module iteration, and why certain modules are intrinsically iterative, could enable biosynthetic engineering approaches to rational skeletal alteration of *trans*-AT PKS products.

