## supplementary information for "Antibiotic skeletal diversification via differential enoylreductase recruitment and module iteration in *trans*-acyltransferase polyketide synthases"

### Methods

##### Creation and modification of protein overproduction constructs

DNA fragments encoding the proteins of interest were amplified by PCR from *B. gladioli* BCC0238 gDNA using Q5® Hot Start High-Fidelity DNA polymerase (New England Biolabs) and amplimers were separated on a 0.8% agarose gel. The desired DNA fragments were excised from the gel, purified using the GeneJET Gel Extraction Kit (Thermo Fisher Scientific) and inserted into pET151 (Thermo Fisher Scientific), pETSUMO (Thermo Fisher Scientific), or pET 28a(+) (New England Biolabs) following the manufacturer's instructions. The EtnK AT domain overproduction construct (pET24a-*etnK*) was synthesized by Epoch Life Sciences and was designed to incorporate an N-terminal octa-histidine tag. Mutant constructs pET-SUMO-*gbnE*-H198V and pET28a(+)-*gbnD2*_Module 6-S941A were generated using the Q5® Site-Directed Mutagenesis Kit (New England Biolabs). Ligation reactions were used to transform *E. coli* TOP10 cells and single colonies were picked and grown overnight at 37 °C and 180 rpm in LB medium containing 50 mg/mL kanamycin or 100 mg/mL ampicillin, as appropriate. Plasmids were isolated from overnight culture using the GeneJET Plasmid Miniprep Kit (Thermo Fisher Scientific) and verified by sequencing. The primers used to amplify an mutagenize protein-encoding DNA fragments are listed in Table S1.

**Table S1: Primers used for amplification and mutagenesis of DNA fragments encoding recombinant proteins used in the study.**

| Construct | Primer |
|---|---|
| pET151-*gbnD1*_ER-ACP | For: *CACC*CCGGAACCGGCTGAGGCG |
| | Rev: *TCA*CGGCGCATGCGTCTCGCT |
| pET151-*gbnD1*_ER | For: *CACC*GCTGCGCCTTCGCCTTCC |
| | Rev: *TCA*CGGCGCATGCGTCTCGCT |
| pET151-*gbnD2*_ACP5 | For: *CACC*CCGAATCCAGCGTCGCTTTCG |
| | Rev: *TCA*GACCTCCGCCTTGGCCGTCTC |
| pET28a(+)-*gbnD1*_ACP3 | For: *ATAT*CATATGACCGCCACGGTTCCGACC |
| | Rev: *ATAT*GAATTCTCAGATCGCGGGCCCGGAGG |
| pET28a(+)-*gbnD2*_ACP4 | For: *ATAT*CATATGCCGGCAACGCAACGCGCC |
| | Rev: *ATAT*GAATTCTCACGCGCTCGCTGTCGCAGG |
| pET28a(+)-*gbnD4*_ACP10 | For: *ATAT*CATATGGCCCAGGCGAAGCCGGTT |
| | Rev: *ATAT*AAGCTTTCAGCCCTCGGGCAGGCCAA |
| pET28a(+)-*gbnD2*_Module 5 | For: *ATAT*GCTAGCGCGGGCGCGAGGACGATG |
| | Rev: *ATAT*AAGCTTTCAGACCTCCGCCTTGGCCGT |
| pET28a(+)-*gbnD2*_Module 5-C209A | For: GGACACCATGgcgTCGTCGTCGCTGACGG |
| | Rev: ATCGCGATGCTGGGACCG |
| pET28a(+)-*gbnD2*_Module 6 | For: *ATA*GCTAGCGACGGCGAGACGGCCAAG |

| | |
|---|---|
| | Rev: *ATA*AAGCTTTCACGGGGCGTTTCCGGTCTT |
| pET28a(+)-*gbnD2*_Module 6-S941A | For: ACTGGGCATCGAT*g*CGGTGGTCG |
| | Rev: TCGGCGAACGGCGTGTCG |
| pET-SUMO-*gbnE*-H198V | For: TTCGGGCGGT*gtT*ACCGATCGC |
| | Rev: TCGGCCTCCACGCAGATGTC |
| pET-SUMO-*gbnE* | For: ATGGCAATGATTACCGCA |
| | Rev: CTATTGCCTCTGGAGTTGAAGTGA |
| pET28a-MBP-TEV-*etnL* | For: CGCGGATCCATGATTACCGCCAAAAGTCT |
| (codon optimized) | Rev: CCGCTCGAGTTAACTGCTAACGCTCTGTG |
| pET28a-*etnE*_ACP5 | For: ATATCATATGAGTCCCGGTGATCTGGCG |
| (codon optimized) | Rev: ATATAAGCTTTCAGGCGCTCACGGCGGTGGC |
| pET28a-*etnG*_ACP10 | For: ATATCATATGACGCCGGCGAGCGAGCAG |
| | Rev: ATATGAATTCTCAGGCCGCGACCTCCGCTCG |

**Protein overproduction and purification**

Single colonies of *E. coli* BL21 (DE3) transformants containing the expression constructs were picked and grown in 10 mL LB containing 50 mg/mL kanamycin or 100 mg/mL ampicillin, as appropriate, at 37 °C and 180 rpm overnight. 1L LB containing 50 mg/mL kanamycin or 100 mg/mL ampicillin, as appropriate, was separately inoculated with each 10 mL overnight culture and grown at 37 °C and 180 rpm until an optical density at 600 nm of 0.6 was reached. 0.5 mM IPTG was added to each culture and growth was continued at 15 °C and 180 rpm for 18-20 h. The cells from each culture were harvested by centrifugation (4,000 *g* at 4 °C for 15 min), separately resuspended in 10 mL of loading buffer (20 mM Tris, 300 mM NaCl, 20 mM imidazole, pH 8.0), and lysed using a cell disruptor (Constant Systems). Each lysate was centrifuged (17,000 *g* at 4 °C for 45 min) and the supernatant was filtered and separately loaded onto a HiTrap Chelating Column (GE HealthCare) that had been pre-loaded with 0.1 M $NiSO_4$ followed by equilibration with loading buffer (20 mM Tris, 300 mM NaCl, 20 mM imidazole, pH 8.0). The loaded columns were washed with 15 mL loading buffer (20 mM Tris, 300 mM NaCl, 20 mM imidazole, pH 8.0) and the proteins were eluted in a stepwise manner using loading buffer containing increasing concentrations of imidazole: 50 mM (5 ml), 100 mM (3 ml), 200 mM (3 ml) and 300 mM (3ml). The presence of the proteins of interest was confirmed by SDS-PAGE and each was further purified by gel filtration using a Superdex 200pg column (GE Healthcare), as required. Fractions containing the same purified protein were combined, exchanged into storage buffer (20 mM Tris, 300 mM NaCl, 10% glycerol, pH 7.4), concentrated using a VivaSpin centrifuge filter (GE Healthcare) with an appropriate molecular weight cut off (MWCO), snap-frozen in liquid nitrogen and stored at -80 °C.

***In vitro* assays:**

**Enoyl reduction assay for ACP domain bound substrate**

The *apo*-ACP domains excised from modules 3, 5, 10 and 12 or contained within the module 1 ACP-ER didomain construct were loaded with crotonyl-Ppant by incubating 100 μM of each protein with 2μM Sfp (purified recombinant PPtase from *Bacillus subtilus*),[1] 10mM $MgCl_2$ and 800 μM crotonyl-CoA in storage buffer (20 mM Tris, 300 mM NaCl, 10% glycerol, pH 7.4) in a total volume of 50 μl. After 30 min incubation at room temperature, UHPLC-ESI-Q-TOF-MS was used to confirm each loading reaction was complete.

For the module 1 ACP-ER didomain enoyl reduction assay, 50 μM crotonylated-*holo*-ACP-ER didomain was incubated with 500 μM NADPH in storage buffer in a total volume of 50 μl. The control reaction was conducted under the same conditions except NADPH was omitted.

For the enoyl reduction assays employing the excised ACP domains, 50 μM of the requisite crotonylated-*holo*-ACP domain was incubated with 50 μM of the module 1 ER domain, GbnE or GbnE-H198V and 500 μM NADPH in storage buffer (50 μl total volume). Control reactions were conducted under the same conditions lacking the module 1 ER domain, GbnE or GbnE-H198V. The reactions were incubated at room temperature for 5h and the nature of the acyl group attached to each ACP domain was analyzed using UHPLC-ESI-Q-TOF-MS/MS (Ppant ejection).

**Module 5 iteration assays**

***KS domain acylation***

The 2,4-hexadienoyl unit was attached to the KS domain by incubating 50 μM module 5 with 1 mM 2,4-hexadienoyl NAC thioester **7** in storage buffer (20 mM Tris, 300 mM NaCl, 10% glycerol, pH 7.4) in a total volume of 50 μl. After 3 h incubation at room temperature, the protein was concentrated to 150 μM using a VivaSpin concentrator with a MWCO of 50 kDa (GE Healthcare). In the control reaction, the mutant construct module 5-C209A was used instead of the wild type protein.

***Iteration assay with intact module 5***

50 μM hexadienoylated-module 5 was incubated with 1 μM Sfp,[1] 10 mM $MgCl_2$, 400 μM malonyl-CoA, 25 μM EtnK AT domain and 800 μM NADPH in the presence / absence of 0.5-50 μM GbnE/GbnE-H198V in storage buffer (20 mM Tris, 300 mM NaCl, 10% glycerol, pH 7.4) in a total volume of 50 μl at room temperature for 20 h. The assay was monitored via UHPLC-ESI-Q-TOF-MS analysis of the intact protein and products were cleaved from the ACP domain via addition of cysteamine to a final concentration of 0.2 M. After 20 h incubation at 4 °C, the mixture was extracted with EtOAc (2 x 1 mL) and the combined extracts were evaporated to

dryness. The residue was dissolved in 100 µL of MeOH and analyzed by UHPLC-ESI-Q-TOF-MS. The instrumentation, elution conditions and settings used for these analyses were the same as those used for analysis of the metabolites produced by *B. gladioli* wild type and in-frame deletion mutants (see below).

***Iteration assay employing module 5 ACP domain* in trans**

The excised module 5 ACP domain was loaded with malonyl-Ppant by incubating 200 µM protein with 2µM Sfp,[1] 10 mM $MgCl_2$ and 800 µM malonyl-CoA in storage buffer in final volume of 50 µl for 30 min at room temperature and UHPLC-ESI-Q-TOF-MS was used to confirm the reaction had gone to completion. 50 µM hexadienoylated-module 5 was incubated with 100 µM malonyated-module 5 *holo*-ACP domain and 800 µM NADPH in the presence / absence of 0.5-50 µM GbnE/GbnE-H198V in storage buffer (final volume 50 µl) at room temperature for 20 h. The species attached to the excised module 5 ACP domain after 2, 5 and 20 h were assessed using UHPLC-ESI-Q-TOF-MS analyses of intact proteins and cysteamine cleavage adducts (as described in the section above on the iteration assay with intact module 5).

**Assay of transacylation from the module 5 ACP domain to the module 6 KS domain**

The module 5 ACP domain was loaded with 2,4-hexadienoyl-Ppant or 4-hexaenoyl-Ppant by incubating 100 µM protein with 2 µM CoaA, 2 µM CoaC, 2 µM CoaD, 2 µM Sfp,[2] 10 mM $MgCl_2$, 5 mM ATP and 1 mM 2,4-hexadienoyl pantetheine thioester **16** or 4-hexenoyl pantetheine thioester **18** in storage buffer (20 mM Tris, 300 mM NaCl, 10% glycerol, pH 7.4) in a total volume of 50 µl for 1.5 h at room temperature. UHPLC-ESI-Q-TOF-MS was used to confirm the reactions had gone to completion. The excess pantetheine thioesters were removed by two rounds of 10-fold dilution with storage buffer and concentration using a VivaSpin centrifuge filter with a MWCO of 5 kDa (GE Healthcare). 100 µM of the hexadienoylated or hexenoylated-module 5 *holo*-ACP domain was incubated with 300 µM of the module 6 KS-ACP-ACP-S941A tridomain in storage buffer (total volume 50 µl) for 8h at room temperature. The acylation state of the module 5 ACP domain after 2 h, 4 h, 6 h and 8 h was assessed by UHPLC-ESI-Q-TOF-MS.

**Module 6 elongation assays**

The KS domain of the module 6 KS-ACP-ACP(S941A) tri-domain was acylated with a 2,4-hexadienoyl or a 4-hexenoyl unit by incubating 50 µM tri-domain with 1mM 2,4-hexadienoyl NAC thioester **7** or 4-hexenoyl NAC thioester **18**, respectively, in storage buffer (20 mM Tris, 300 mM NaCl, 10% glycerol, pH 7.4) in a total volume of 50 µl at room temperature for 3 h. Acylation of the tri-domain was confirmed by UHPLC-ESI-Q-TOF-MS analysis. The acylated tri-domains were incubated with 1 µM Sfp,[1] 500 µM malonyl-CoA and 10mM $MgCl_2$ in storage buffer (total volume of 50 µl). Control reactions were conducted under the same conditions except that the 2,4-hexadienoyl and 4-hexenoyl NAC thioesters were omitted. After 6 h incubation at room temperature the acylation state of the tri-domains was assessed using UHPLC-ESI-Q-TOF-MS.

**Intact protein analysis using UHPLC-ESI-Q-TOF-MS**

Purified recombinant proteins and proteins of interest in enzymatic activity assays were analysed using a Bruker MaXis II ESI-Q-TOF-MS connected to a Dionex 3000 RS UHPLC fitted with an ACE C4-300 RP column (100 x 2.1 mm, 5 µm, 30 °C). The column was eluted with a linear gradient of 5%-100% MeCN containing 0.1% formic acid at a flow rate of 0.2 ml/min for 30 min. The mass spectrometer was operated in positive ion mode with a scan range of 200–3000 *m/z*. Source conditions were: end plate offset at −500 V; capillary at −4500 V; nebulizer gas ($N_2$) at 1.8 bar; dry gas ($N_2$) at 9.0 L $min^{-1}$; dry temperature at 200 °C. Ion transfer conditions were: ion funnel RF at 400 Vpp; multiple RF at 200 Vpp; quadrupole low mass at 300 *m/z*; collision energy at 8.0 eV; collision RF at 2000 Vpp; transfer time at 110.0 µs; pre-pulse storage time at 10.0 µs.

**Construction and complementation of *in-frame* gene deletion mutants**

In-frame gene deletions were constructed in *Burkholderia gladioli* BCC1622 (because the original gladiolin producer *B. gladioli* BCC0238 was not suitable for genetic manipulation using this system).[3] The primers listed in Table S2 were used to amplify the 5' and 3' regions flanking each gene. The resulting amplimers were cloned into pGPI-SceI and transferred into *E. coli* SY327 by electroporation. The integrity of the resulting plasmids was confirmed by sequencing and they were mobilized into *Burkholderia gladioli* BCC1622 by tri-parental mating.[4] Exconjugants were selected for using 150 µg/ml trimethoprim and 600 U/ml polymyxin B and subjected to PCR screening to identify single crossover mutants using the primers listed in Table S2. A confirmed single crossover mutant for each gene of interest was transformed with pDAI-SceI by tri-parental mating. Exconjugants were selected for using 200 µg/ml tetracycline and 600 U/ml polymyxin B and subjected to PCR screening to identify double crossover mutants using the primers listed in Table S2. The resulting mutants were confirmed as trimethoprim sensitive. Finally, the pDAI-SceI plasmid was removed from one of the double crossover mutants by growing it on M9 minimal medium containing 15% sucrose. Single colonies were picked and confirmed to be sensitive to tetracycline.

For in *trans* complementation of deletion mutants, the gene of interest was amplified using the primers listed in Table S2. The resulting amplimers were cloned into pMLBAD and used to transform *E. coli* Top10 by electroporation. The integrity of the resulting plasmids was confirmed by sequencing and they were mobilized into the appropriate in-frame deletion mutant of *Burkholderia gladioli* BCC1622 by tri-parental mating as described above.[4] Exconjugants were selected for using 150 μg/ml trimethoprim and 600 U/ml polymyxin B and confirmed by $Tp^R$ phenotype and PCR with the primer pairs used for initial amplification of each gene.

**Table S2: Primers for gene in-frame deletion and complementation**

| Construct | Primer (restriction site) |
|---|---|
| pGPI-*gbnE* | 5'-For: ATATCTAGAAGCAAGCGGTCCGACAGG (*Xba*I) |
| | 5'-Rev: ATAAAGCTTATCGGCGTTGCCAAGCGA (*Hin*dIII) |
| | 3'-For: TATAAGCTTTCGATGGCCGATTCGGGC (*Hin*dIII) |
| | 3'-Rev: ATAGAATTCTGGCGGGTCGAGCCCATC (*Eco*RI) |
| | Screening-For: CTGTTGCAAGCGCACGATTAGC |
| | Screening-Rev: GACCAGGGGCAGTCGCAGCGA |
| pGPI-*gdsB_ER* | 5'-For: *gatcccaagcttcttctaga*GCACTGTTCGAGAAGTAC |
| | 5'-Rev: *gttgggtgaa*GTAGGCGTACTTGAGCTT |
| | 3'-For: *gtacgcctac*TTCACCCAACTGCTCCG |
| | 3'-Rev: *aacggctgacatgggaattc*GAAGATCGACGAGAG |
| | Screening-For: CAGGGCGTCGAGACATTCC |
| | Screening_Rev: CCTTGTCGCGCACGGATT |
| pMLBAD-*gbnE* | For: *tgggctagcaggaggaattc*ATGGCAATGATTACCGCA |
| | Rev: *tgggctagcaggaggaattc*ATGGCAATGATTACCGCA |
| pMLBAD_*etnL* | For: CCGGAATTCATGATTACCGCCAAAAGTCT (*Eco*RI) |
| (codon optimized) | Rev: CCCAAGCTTTTAACTGCTAACGCTCTGTG (*Hin*dIII) |

**Metabolite production, extraction, and UHPLC-ESI-Q-TOF-MS analysis**

*Burkholderia* strains were grown on basal salts agar medium containing 0.4% (w/v) glycerol (BSM-G) at 30 °C for 72 h. For strains harboring pMLBAD-derived plasmids, 50 μg/ml trimethoprim and 0.2% (w/v) L-arabinose were added to the medium. Following incubation, the biomass was scraped off the plate, the agar was cut into 0.5 cm cubes and extracted twice using MeCN and the extract was passed through a 0.2 μm nylon centrifuge filter (Thermo Scientific). The filtered extract was analyzed using a Dionex UltiMate 3000 UHPLC connected to a Zorbax Eclipse Plus column (C18, 100 × 2.1 mm, 1.8 μm) coupled to a Bruker MaXis IMPACT ESI-Q-TOF mass spectrometer. The column was eluted with a gradient of 20% to 100% MeCN containing 0.1% formic acid at flow rate of 0.2 ml/min over 35 min. The mass spectrometer was operated in positive ion mode with a scan range of 50-3000 *m/z*. Source conditions were: end plate offset at −500 V; capillary at −4500 V; nebulizer gas ($N_2$) at 1.6 bar; dry gas ($N_2$) at 8 L $min^{-1}$; dry temperature at 180 °C. Ion transfer conditions were: ion funnel RF at 200 Vpp; multiple RF at 200 Vpp; quadrupole low mass at 55 *m/z*; collision energy at 5.0 eV; collision RF at 600 Vpp; ion cooler RF at 50–350 Vpp; transfer time at 121 s; pre-pulse storage time at 1 s. Calibration was performed with 1 mM sodium formate through a loop injection of 20 µL at the start of each run.

**Synthesis of (2*E*, 4*E*)-2,4-hexadienoyl NAC thioester 7 and pantetheine thioester 17**

***(2*E*, 4*E*)-2,4-hexadienoyl NAC thioester 7***

(2*E*, 4*E*)-2,4-hexadienoyl NAC thioester **7** was synthesized using the same procedure as that used for the synthesis of (*E*)-4-hexenoyl NAC thioester **19** (see below) except that sorbic acid (100 mg, 0.89 mmol) was used in place of (*E*)-4-hexenoic acid and the scale of the reaction was adjusted accordingly. This afforded the desired product as a white solid (176 mg, 93 %).

$\nu_{max}/cm^{-1}$ (neat) 3327 (NH), 1665, 1532 (C=O); $\delta_H$ (500 MHz; $CDCl_3$) 7.20 (1H, dd, J 15.0 and 10.5, C*H*CHCOS), 6.25 (1H, dq, J 15.0 and 7.0, $CH_3$C*H*), 6.16 (1H, ddd, J 15.0, 11.0 and 1.0, $CH_3$CHC*H*), 6.07 (1H, d, J 15.0, C*H*COS), 5.91 (1H, m, N*H*), 3.45 (2H, q, J 6.0, C*H*$_2$NH), 3.07 (2H, t, J 6.5, C*H*$_2$S), 1.96 (3H, s, COC*H*$_3$), 1.85 (3H, d, J 6.5, CHC*H*$_3$); $\delta_C$ (125 MHz, $CDCl_3$) 191.6 (*C*OS), 170.3 (*C*ON), 142.1 ($CH_3$*C*H), 142.0 (*C*HCHCOS), 129.7 ($CH_3$CH*C*H), 125.8 (CH*C*HCOS), 40.0 (*C*H$_2$NH), 28.5 (*C*H$_2$S), 23.4 (CO*C*H$_3$), 14.6 (CH*C*H$_3$); HRMS (ESI) cald. for $C_{10}H_{15}NNaO_2S$ (M + $Na^+$) requires 236.0721, found 236.0720.

***(2*E*, 4*E*)-2,4-hexadienoyl pantetheine thioester 17***

(2*E*, 4*E*)-2,4-hexadienoyl pantetheine thioester **17** was synthesized as described previously.[5]

**Synthesis of (*E*)-4-hexenoyl pantetheine thioester 18 and NAC thioester 19**

***(*E*)-4-hexenoyl pantetheine thioester* 18**

(*R*)-*S*-(2-(3-(2,2,5,5-tetramethyl-1,3-dioxane-4-carboxamido)propanamido)ethyl) (*E*)-hex-4-enethioate was synthesized using the same procedure as that used for the synthesis of (*E*)-4-hexenoyl NAC thioester **19** (see below) except that (*R*)-*N*-(3-((2-mercaptoethyl)amino)-3-oxopropyl)-2,2,5,5-tetramethyl-1,3-dioxane-4-carboxamide[5] (134 mg, 0.42 mmol, 1.4 equiv.) was used in place of *N*-(2-mercaptoethyl)acetamide and the scale of the reaction was adjusted accordingly. This afforded *(R)*-*S*-(2-(3-(2,2,5,5-tetramethyl-1,3-dioxane-4-carboxamido)propanamido)ethyl) *(E)*-hex-4-enethioate as a colorless oil (123 mg, 76 %).

$\nu_{max}$/cm$^{-1}$ (neat) 3338 (NH), 1665, 1538 (C=O); $\delta_H$ (500 MHz; $CDCl_3$) 7.02 (1H, br. t, J 6.0, N*H*), 6.16 (1H, br. t, J 5.5, N*H*$CH_2CH_2S$), 5.48 (1H, dq, J 15.5 and 6.5, C*H*$CH_3$), 5.37 (1H, dt, J 15.5 and 6.5, C*H*$CHCH_3$), 4.07 (1H, s, C*H*CONH), 3.67 (1H, d J 11.5, $CH_2OC(CH_3)_2$), 3.60-3.35 (4H, m, NHC$H_2$, C$H_2CH_2S$), 3.27 (1H, d J 12.0, C$H_2OC(CH_3)_2$), 3.00 (2H, t, J 6.5, C$H_2$S), 2.61 (2H, t, J 7.5, C$H_2$COS), 2.41 (2H, t, J 6.0, C$H_2$CONH), 2.32 (2H, q, J 7.0, C$H_2CH_2$COS), 1.63 (3H, d, J 6.0, CHC$H_3$), 1.45 (3H, s, OC(C$H_3$)$_2$), 1.41 (3H, s, OC(C$H_3$)$_2$), 1.03 (3H, s, $CH_2C(CH_3)_2$), 0.96 (3H, s, $CH_2C(CH_3)_2$); $\delta_C$ (125 MHz, $CDCl_3$) 199.3 (*C*$O_2$S), 171.3 ($CH_2$*C*ONH), 170.2 (CH*C*ONH), 128.6 ($CH_3$CH*C*H), 126.8 ($CH_3$*C*H), 99.2 (O*C*($CH_3$)$_2$), 77.3 (*C*H), 71.6 (*C*$H_2OC(CH_3)_2$), 44.1 (*C*$H_2$COS), 39.7 (*C*$H_2CH_2S$), 36.1 (*C*$H_2$CONH), 34.9 (*C*$H_2$NH), 33.1 ($CH_2$*C*($CH_3$)$_2$), 29.6 (OC(*C*$H_3$)$_2$), 28.6 **(***C*$H_2CH_2$COS**)**, 28.5 (*C*$H_2$S), 22.3 ($CH_2C$(*C*$H_3$)$_2$), 19.0 (OC(*C*$H_3$)$_2$), 18.8 ($CH_2C$(*C*$H_3$)$_2$), 18.0 (CH*C*$H_3$); HRMS (ESI) cald. for $C_{20}H_{34}N_2NaO_5S$: 437.2086, found 437.2085 ([M+Na]$^+$); $[\alpha]_D^{27}$ (c 0.1, $CHCl_3$): +25.3.

*(R)*-*S*-(2-(3-(2,2,5,5-tetramethyl-1,3-dioxane-4-carboxamido)propanamido)ethyl) *(E)*-hex-4-enethioate (70 mg, 0.17 mmol, 1.0 equiv.) was stirred in AcOH : $H_2O$ (2 : 1, 3 mL), for 16 h at room temperature. The mixture was concentrated *in vacuo* and purified using silica chromatography ($CH_2Cl_2$ / MeOH, 9:1) to give the product as a colorless oil (52 mg, 82 %).

$\nu_{max}$/cm$^{-1}$ (neat) 3357 (OH), 3301 (NH), 1657, 1541 (C=O); $\delta_H$ (500 MHz; $CD_3OD$) 5.49 (1H, dq, J 15.5 and 6.0, C*H*$CH_3$), 5.41 (1H, dt, J 15.5 and 6.5, C*H*$CHCH_3$), 3.89 (1H, s, C*H*CONH), 3.53-3.37 (6H, m, NHC$H_2$, C$H_2CH_2S$, C$H_2$OH), 2.99 (2H, t, J 6.5, C$H_2$S), 2.62 (2H, t, J 7.5, C$H_2$COS), 2.40 (2H, t, J 6.5, C$H_2$CONH), 2.30 (2H, q, J 7.0, C$H_2CH_2$COS), 1.63 (3H, d, J 6.5, CHC$H_3$), 0.92 (6H, s, C(C$H_3$)$_2$); $\delta_C$ (125 MHz, $CD_3OD$) 200.0 (*C*$O_2$S), 176.1 ($CH_2$*C*ONH), 173.9 (CH*C*ONH), 130.0 ($CH_3$CH*C*H), 127.5 ($CH_3$*C*H), 77.3 (*C*H), 70.4 (*C*$H_2$OH), 44.8 (*C*$H_2$COS), 40.4 ($CH_2$*C*($CH_3$)$_2$), 40.1 (*C*$H_2CH_2S$), 36.4 (*C*$H_2$CONH), 36.3 (*C*$H_2$NH), 29.5 (*C*$H_2CH_2$COS), 29.1 (*C*$H_2$S), 21.3 ($CH_2C$(*C*$H_3$)$_2$), 20.9 ($CH_2C$(*C*$H_3$)$_2$), 18.0 (CH*C*$H_3$); HRMS (ESI) cald. for $C_{17}H_{30}N_2NaO_5S$: 397.1773, found 397.1770 ([M+Na]$^+$); $[\alpha]_D^{27}$ (c 0.2, MeOH): +46.3.

***(*E*)-4-hexenoyl NAC thioester* 19**

To a solution of (*E*)-4-hexenoic acid (76 mg, 0.67 mmol, 1.3 equiv.),[6] *N*-(2-mercaptoethyl)acetamide (88 mg, 0.74 mmol, 1.4 equiv.) and DMAP (18 mg, 0.16 mmol, 0.3 equiv.) in $CH_2Cl_2$ (5 mL), was added EDC (141 mg, 0.75 mmol, 1.4 equiv.) at 0 °C. The resulting mixture was stirred at room temperature overnight, quenched by the addition of 2 M HCl, extracted with $CH_2Cl_2$ (3 x 10 mL), washed with brine (10 mL), dried ($MgSO_4$), filtered and concentrated *in vacuo*. Purification using silica gel chromatography (EtOAc) afforded the desired product as a colorless oil (131 mg, 91 %).

$\nu_{max}$/cm$^{-1}$ (neat) 3326 (NH), 1659, 1545 (C=O); $\delta_H$ (500 MHz; $CDCl_3$) 5.80 (1H, br. s, N*H*), 5.49 (1H, dq, J 15.0 and 6.5, $CH_3$C*H*), 5.38 (1H, dt, J 15.5 and 6.5, $CH_3$CHC*H*), 3.43 (2H, q, J 6.0, C*H*$_2$NH), 3.02 (2H, t, J 6.5, C*H*$_2$S), 2.63 (2H, t, J 7.5, C*H*$_2$COS), 2.34 (2H, q, J 7.0, C*H*$_2$$CH_2$COS), 1.96 (3H, s, COC*H*$_3$), 1.64 (3H, d, J 6.5, CHC*H*$_3$); $\delta_C$ (125 MHz, $CDCl_3$) 199.8 (*C*OS), 170.4 (*C*ON), 128.6 ($CH_3$CH*C*H), 126.9 ($CH_3$*C*H), 44.1 (*C*H$_2$COS), 39.9 (*C*H$_2$NH), 28.6 (*C*H$_2$$CH_2$COS), 28.6 (*C*H$_2$S), 23.4 (CO*C*H$_3$), 18.0 (CH*C*H$_3$); HRMS (ESI) cald. for $C_{10}H_{17}NNaO_2S$ ($[M+Na]^+$) requires 238.0878, found 238.0880.

#### Figures

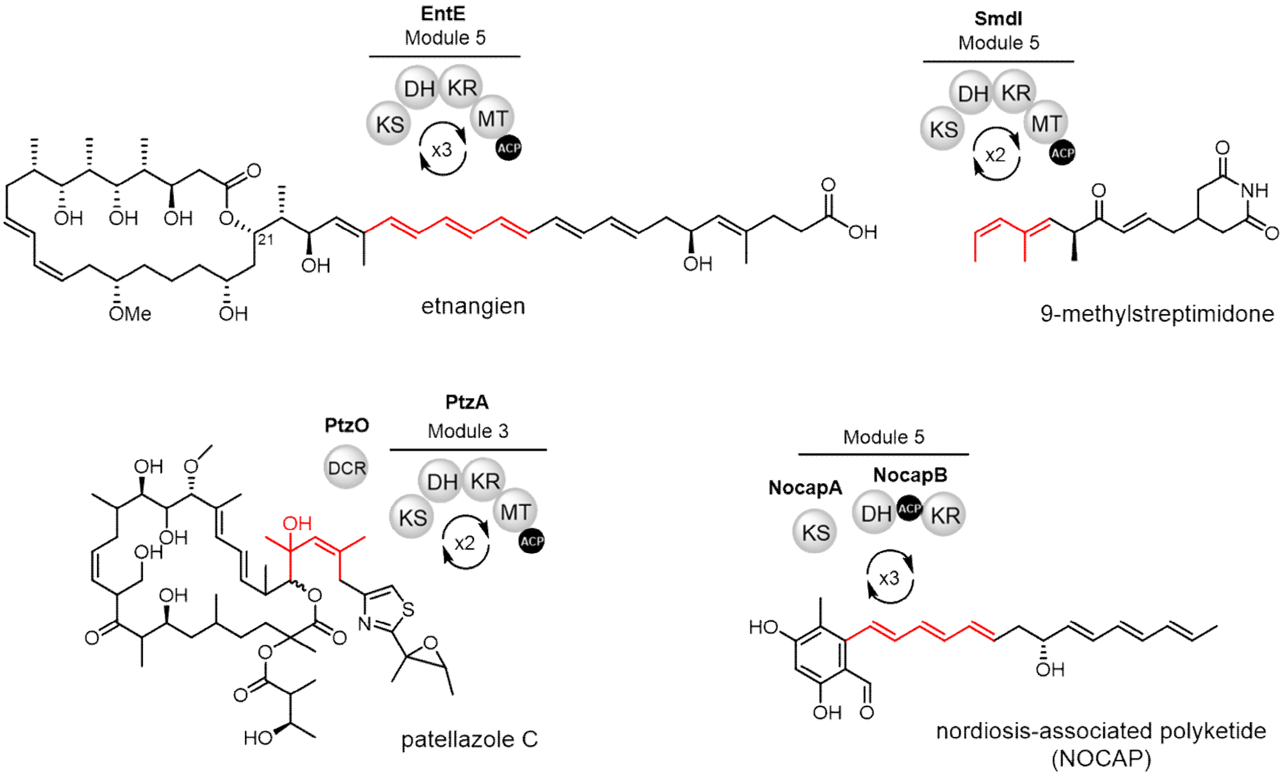


**Figure S1. Examples of polyketides assembled by *trans*-AT PKSs containing putative intrinsically iterative modules.** Moieties derived from module iteration are highlighted in red.

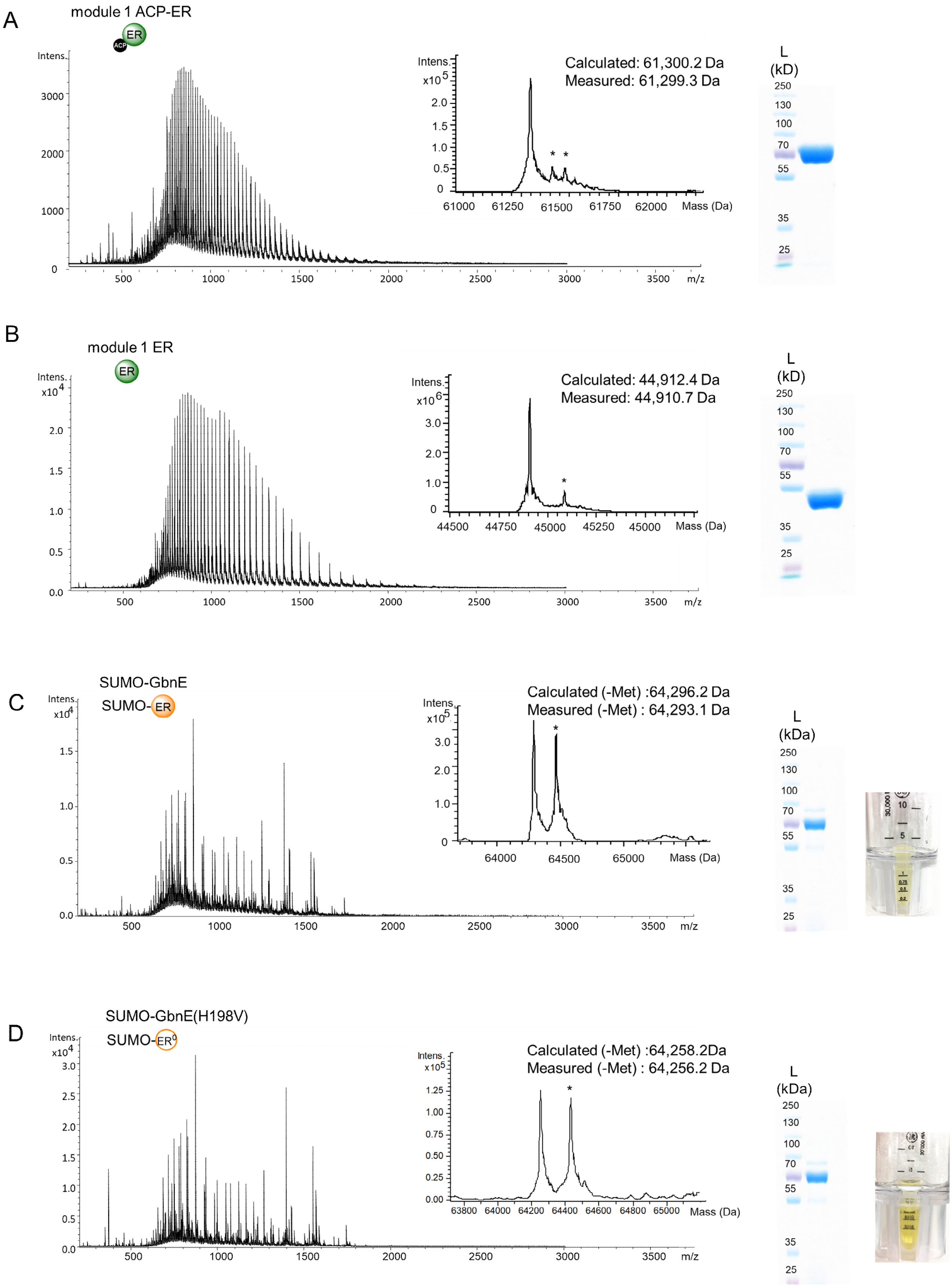
A
module 1 ACP-ER
ACP
ER
Intens.
Intens. x10^5
Calculated: 61,300.2 Da
Measured: 61,299.3 Da
Mass (Da)
m/z
L
(kD)
B
module 1 ER
ER
Intens. x10^4
Intens. x10^6
Calculated: 44,912.4 Da
Measured: 44,910.7 Da
Mass (Da)
m/z
L
(kD)
C
SUMO-GbnE
SUMO-ER
Intens. x10^4
Intens. x10^5
Calculated (-Met) :64,296.2 Da
Measured (-Met) : 64,293.1 Da
Mass (Da)
m/z
L
(kDa)
D
SUMO-GbnE(H198V)
SUMO-ER°
Intens. x10^4
Intens. x10^5
Calculated (-Met) :64,258.2Da
Measured (-Met) : 64,256.2 Da
Mass (Da)
m/z
L
(kDa)

E

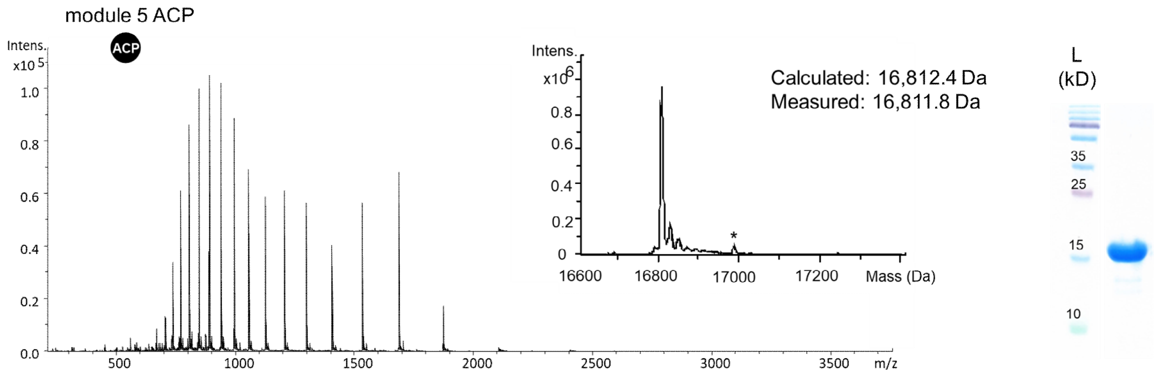
module 5 ACP
ACP
Calculated: 16,812.4 Da
Measured: 16,811.8 Da
Intens.
Mass (Da)
m/z
L
(kD)

F

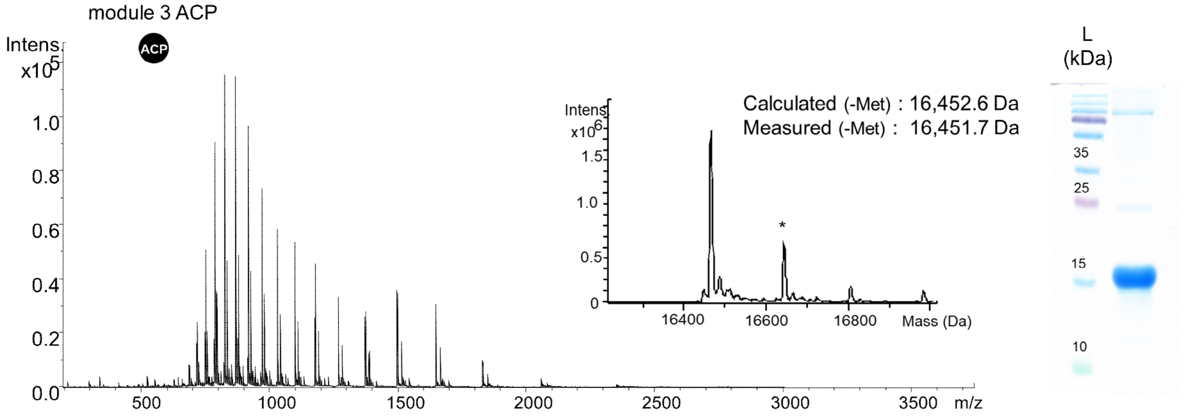
module 3 ACP
ACP
Calculated (-Met) : 16,452.6 Da
Measured (-Met) : 16,451.7 Da
Intens.
Mass (Da)
m/z
L
(kDa)

G

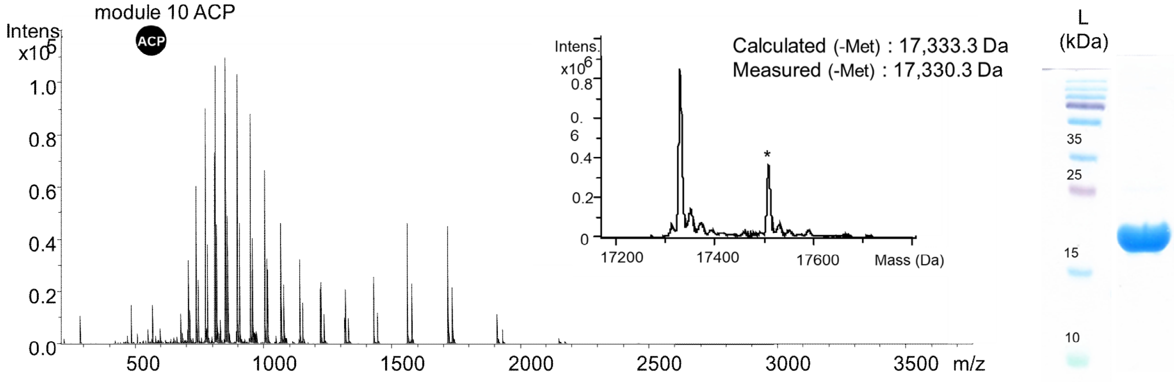
module 10 ACP
ACP
Calculated (-Met) : 17,333.3 Da
Measured (-Met) : 17,330.3 Da
Intens.
Mass (Da)
m/z
L
(kDa)

H

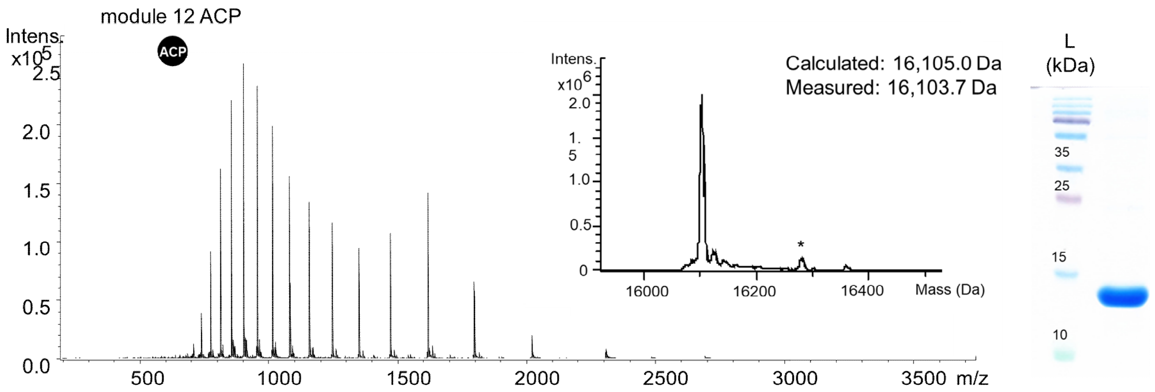
module 12 ACP
ACP
Calculated: 16,105.0 Da
Measured: 16,103.7 Da
Intens.
Mass (Da)
m/z
L
(kDa)

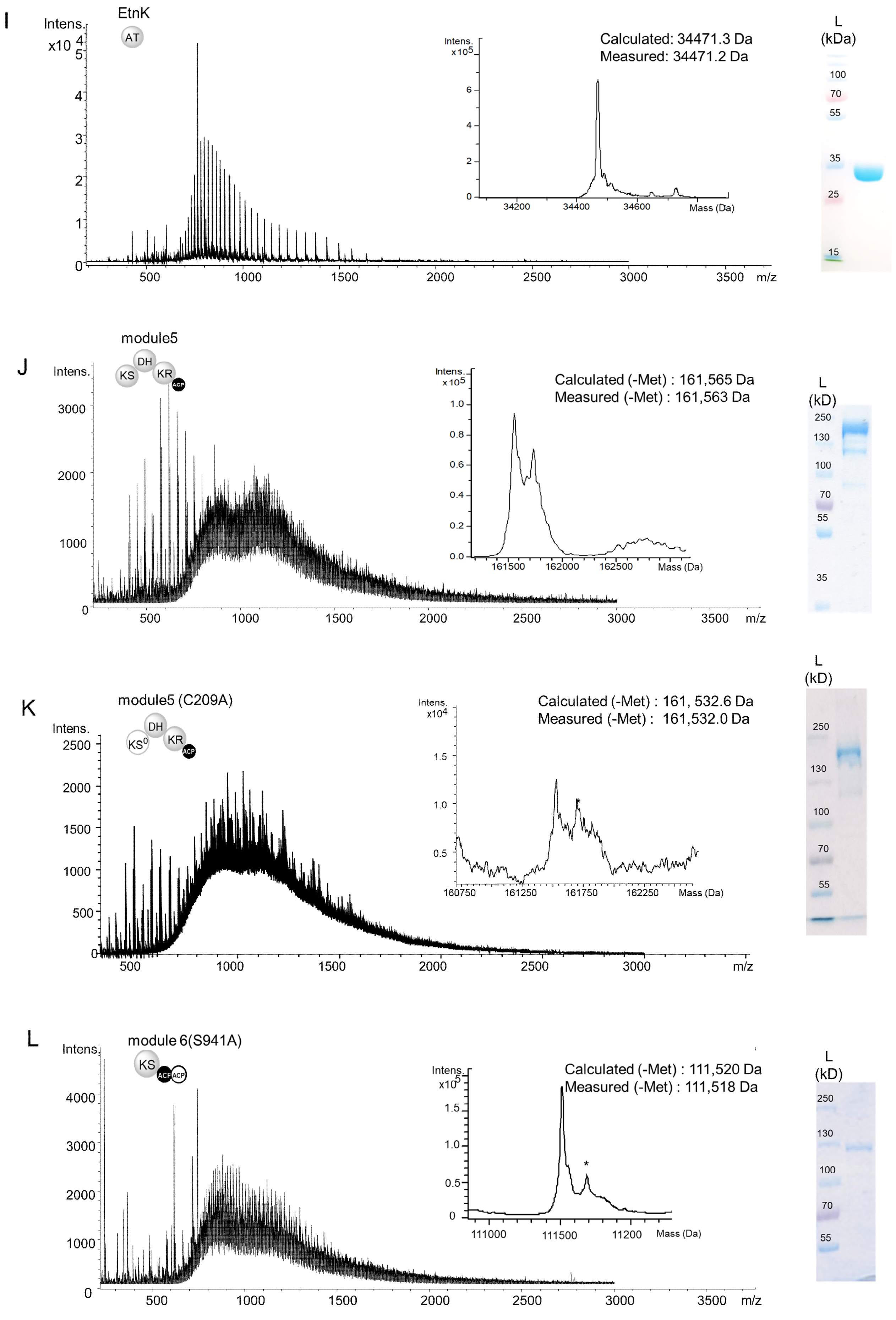
I
EtnK
AT
Intens.
x10^4
m/z
Calculated: 34471.3 Da
Measured: 34471.2 Da
Mass (Da)
L
(kDa)
J
module5
KS
DH
KR
ACP
Calculated (-Met) : 161,565 Da
Measured (-Met) : 161,563 Da
L
(kD)
K
module5 (C209A)
KS0
Calculated (-Met) : 161, 532.6 Da
Measured (-Met) : 161,532.0 Da
L
module 6(S941A)
Calculated (-Met) : 111,520 Da
Measured (-Met) : 111,518 Da

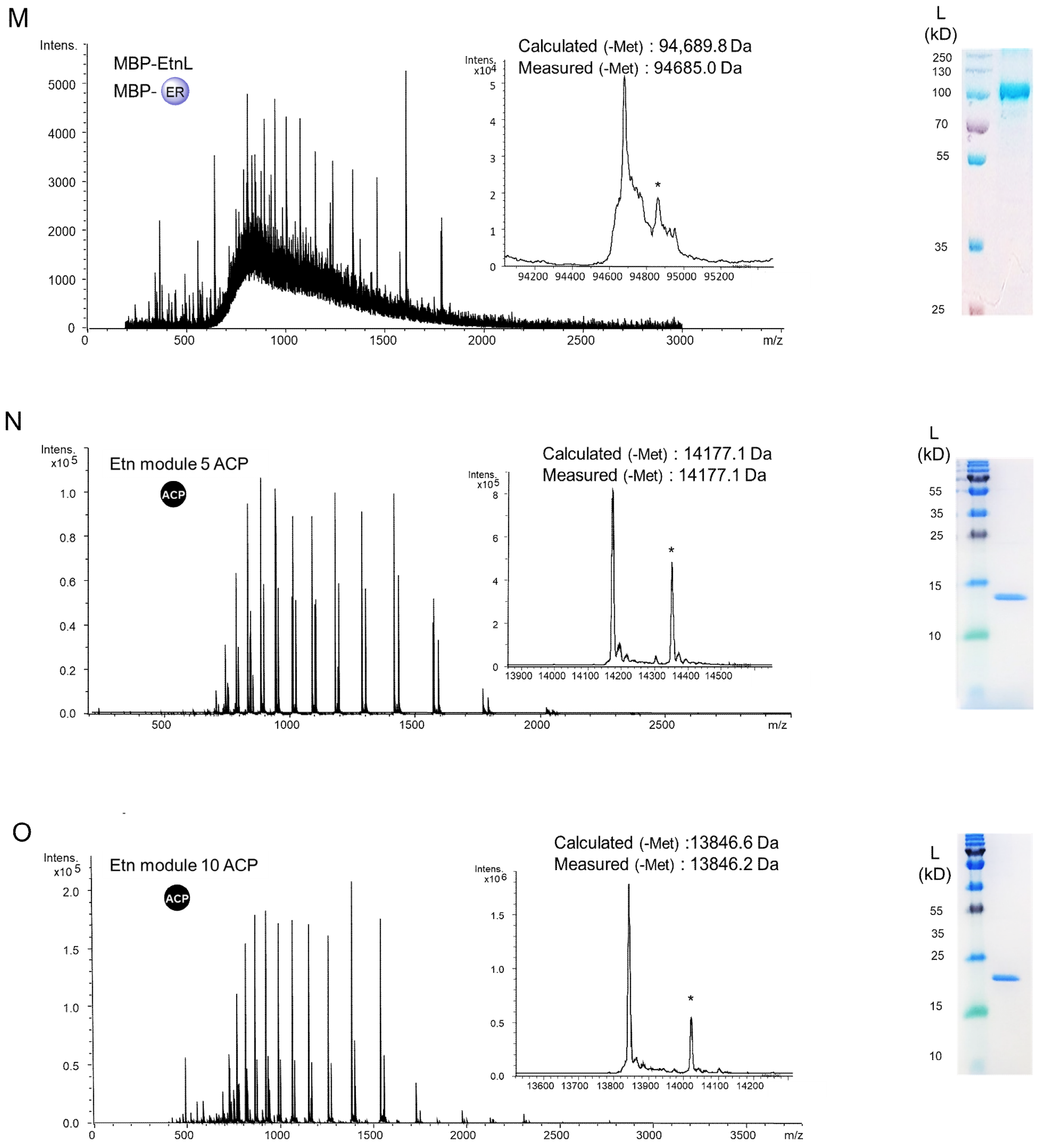


**Figure S2. UHPLC-ESI-Q-TOF-MS and SDS-PAGE characterization of purified recombinant proteins.** Raw mass spectra (left), deconvoluted mass spectra (middle) and SDS-PAGE gels (right) of purified recombinant proteins. Peaks labeled with '*' correspond to a gluconylated form of the protein resulting in a mass increase of 178 Da. Purified SUMO-GbnE, SUMO-GbnE-H198V and MBP-EtnL were yellow, consistent with a bound flavin cofactor. The N-terminal Met residue of some proteins was removed by an internal aminopeptidase during overproduction in *E. coli* BL21 (DE3)[7] -indicated as '(-Met)' in the calculated and measured masses.

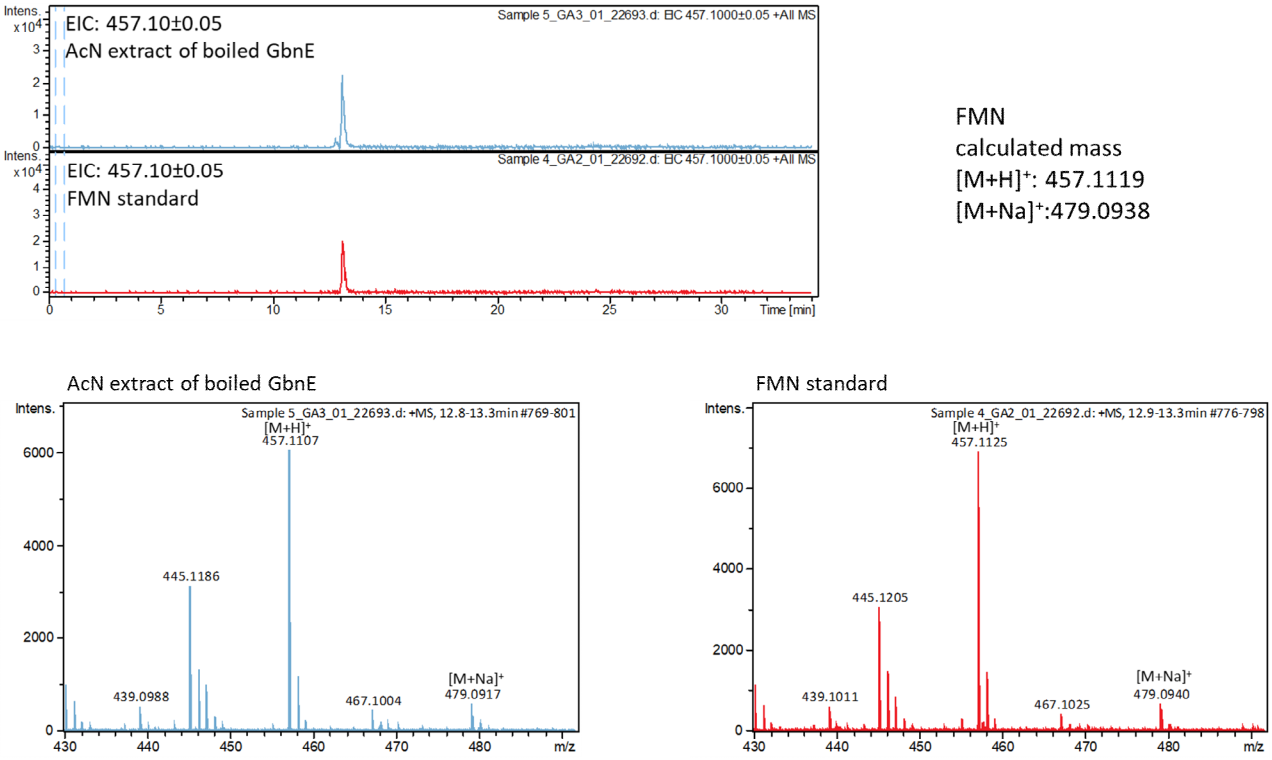


**Figure S3. UHPLC-ESI-Q-TOF-MS analysis of an MeCN extract of GbnE confirming it purifies with FMN bound.** The extracted ion chromatogram (top panel) corresponding to $[M+H]^+$ for FMN (*m/z* = 457.10± 0.05) from an MeCN extract of boiled GbnE (top chromatogram) contained a species with the same retention time as that for an authentic standard of FMN (bottom chromatogram). High resolution mass spectra of FMN in the MeCN extract of GbnE (bottom left panel) and an authentic standard of FMN (bottom right panel). The *m/z* values observed for the $[M+H]^+$ and $[M+Na]^+$ ions are in good agreement with the calculated values (top right).

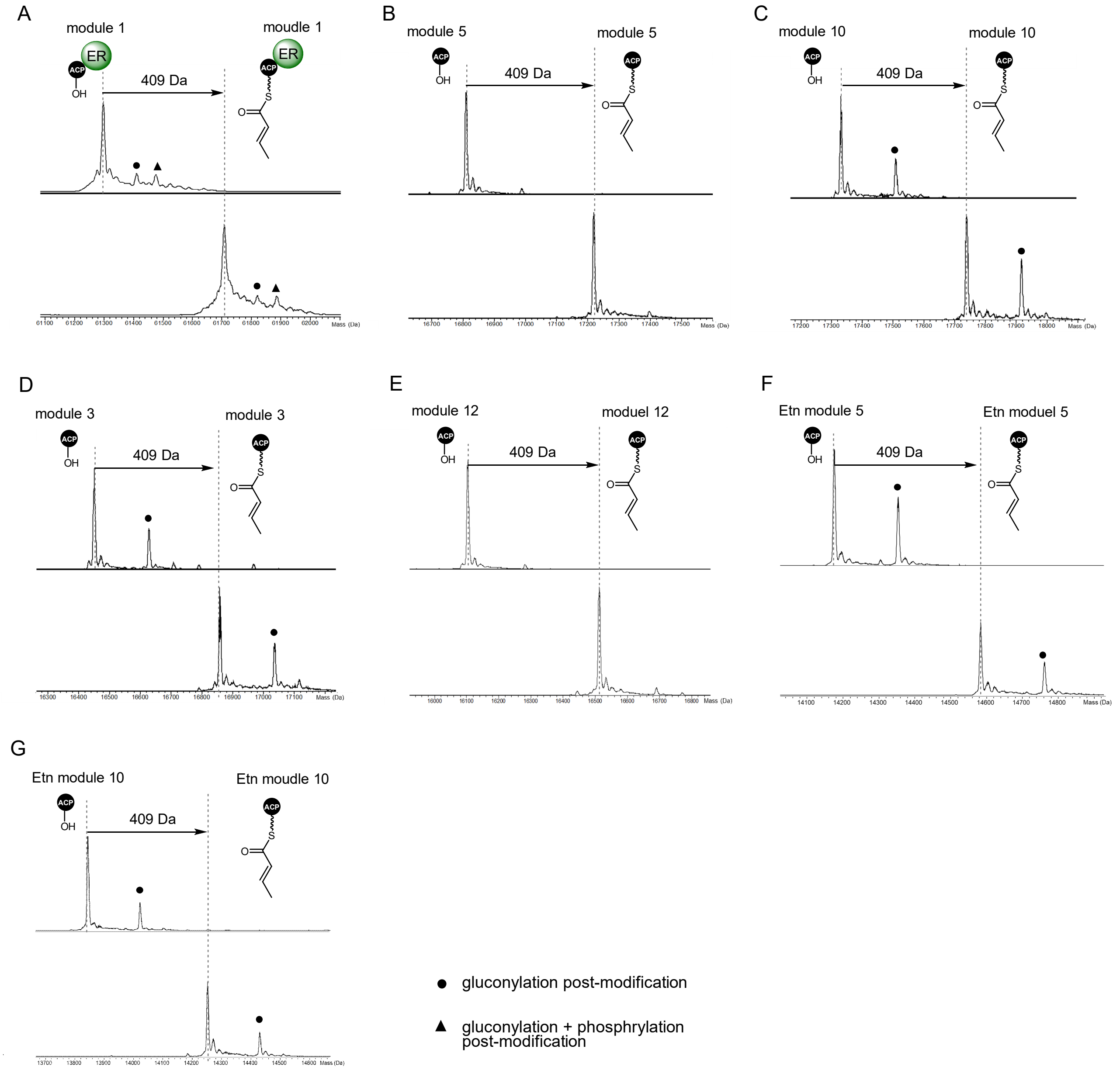


**Figure S4. Deconvoluted mass spectra from UHPLC-ESI-Q-TOF-MS analysis of *apo*-ACP domain conversion to crotonylated *holo* forms.** Gluconylated and gluconylated + phosphorylated species are indicated with a dot and triangle, respectively. A mass shift of 409 Da is observed for conversion from the *apo* to crotonylated *holo* form.

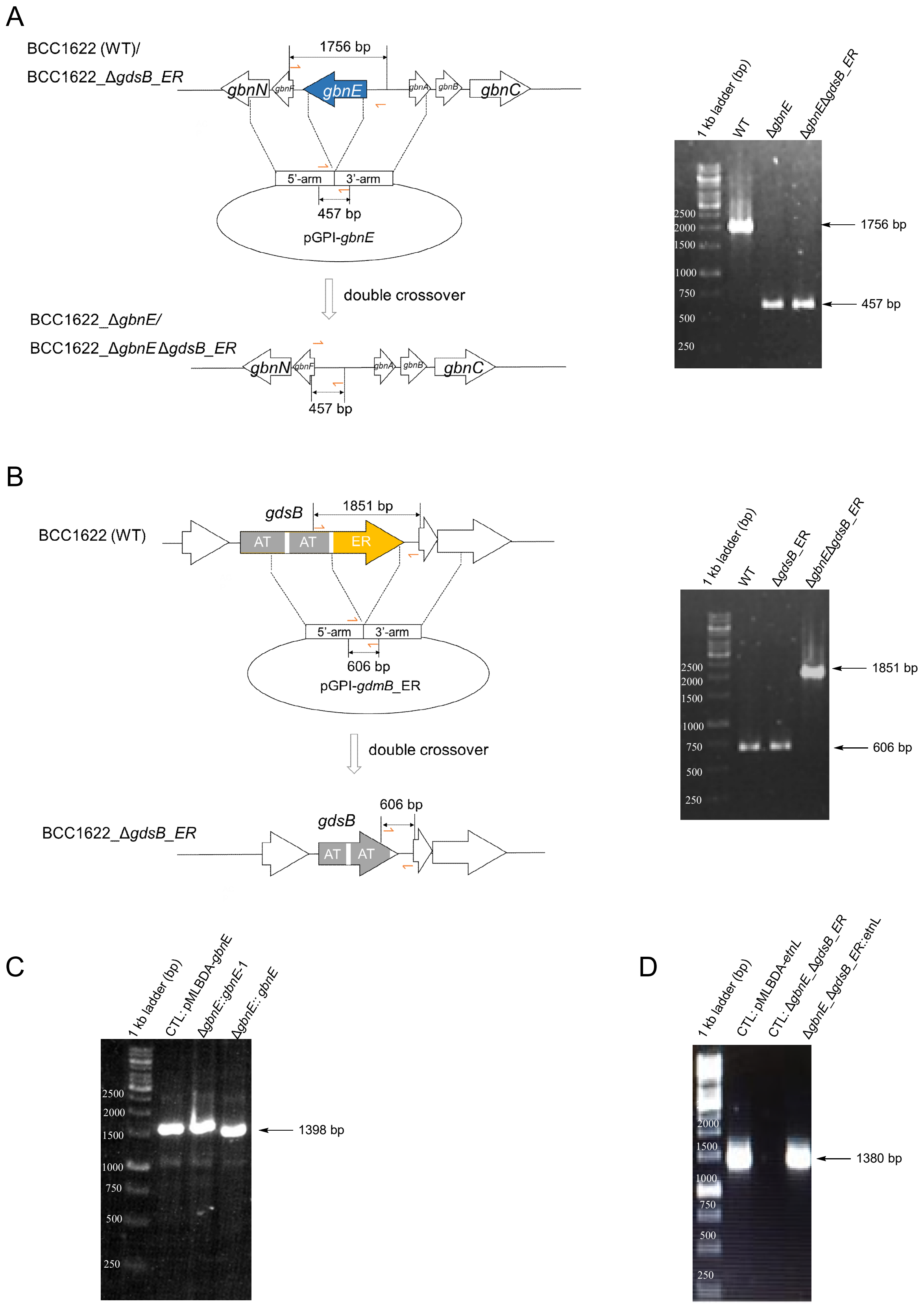


**Figure S5. Construction of in-frame deletion mutants in *B. gladioli* BCC1622.** (A) Scheme for construction of *gbnE* in-frame deletion and PCR confirmation of the expected genotype. (B) Scheme for construction of *gdsB*_ER in-frame deletions and PCR confirmation of the expected genotype. (C) PCR confirmation of the genotype of the Δ*gbnE*::*gbnE* mutant. (D) PCR confirmation of the genotype of the Δ*gdsB*_ERΔ*gbnE*::*etnL*

mutant.

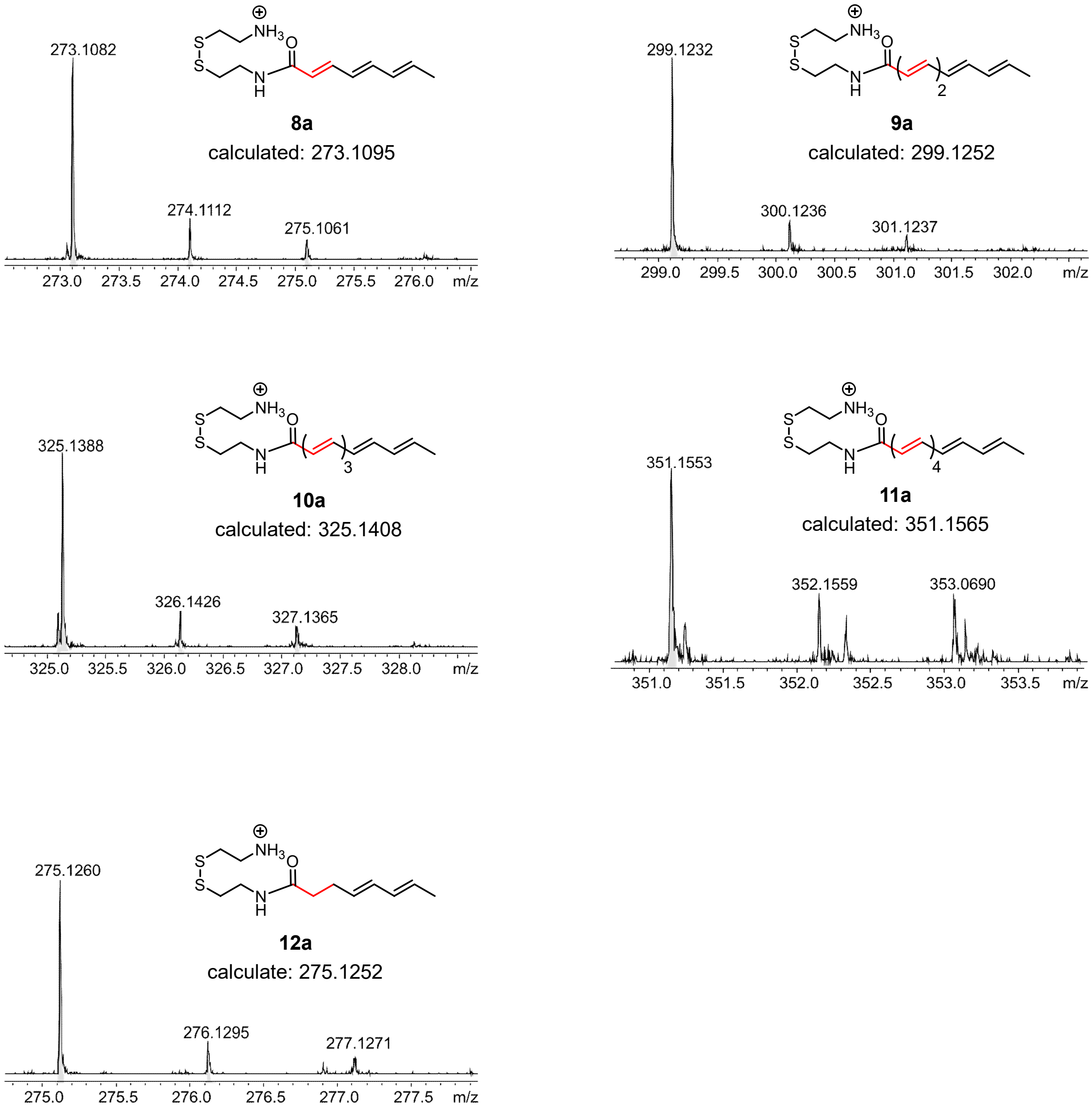


**Figure S6. High resolution mass spectra of cysteamine cleavage adducts observed in UHPLC-ESI-Q-TOF-MS analyses of module 5 *in vitro* reconstitution assays in the presence and absence of GbnE or an H198V mutant.** Representative spectra of the $[M+H]^+$ ions for **8a**, **9a**, **10a**, **11a** and **12a** and comparison with the calculated *m/z* for each species.

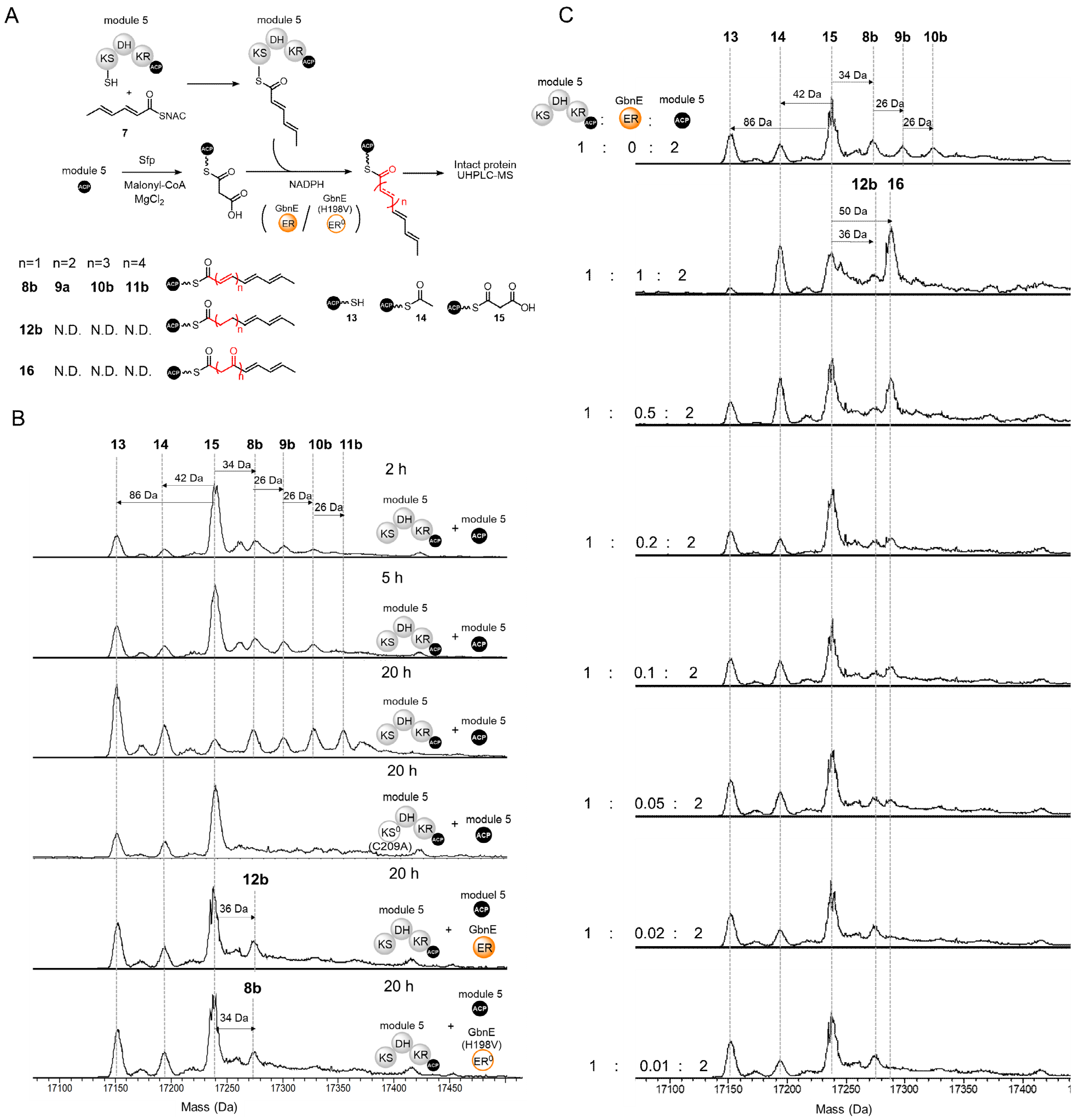

**Figure S7. Deconvoluted mass spectra from UHPLC-ESI-Q-TOF-MS analysis of the module 5 *in vitro* reconstitution in the presence of the excised module 5 *holo*-ACP domain and presence/absence of GbnE/GbnE(H198V).** (A) Scheme illustrating the assay design. (B) Spectra of the excised module 5 ACP domain from the *in vitro* reconstitution assays in the absence of GbnE after 2 h (top), 5 h (second from top) and 20 h (third from top) and in the presence of GbnE after 20 h (second from bottom) and the presence of GbnE(H198V) after 20 h (bottom). The spectrum of the excised module 5 ACP domain from a control reaction in which module 5 was replaced by a C209A mutant with an inactive KS domain is third from bottom. (C) Spectra of the excised module 5 ACP domain from the *in vitro* reconstitution assays containing varying amounts of GbnE relative to module 5.

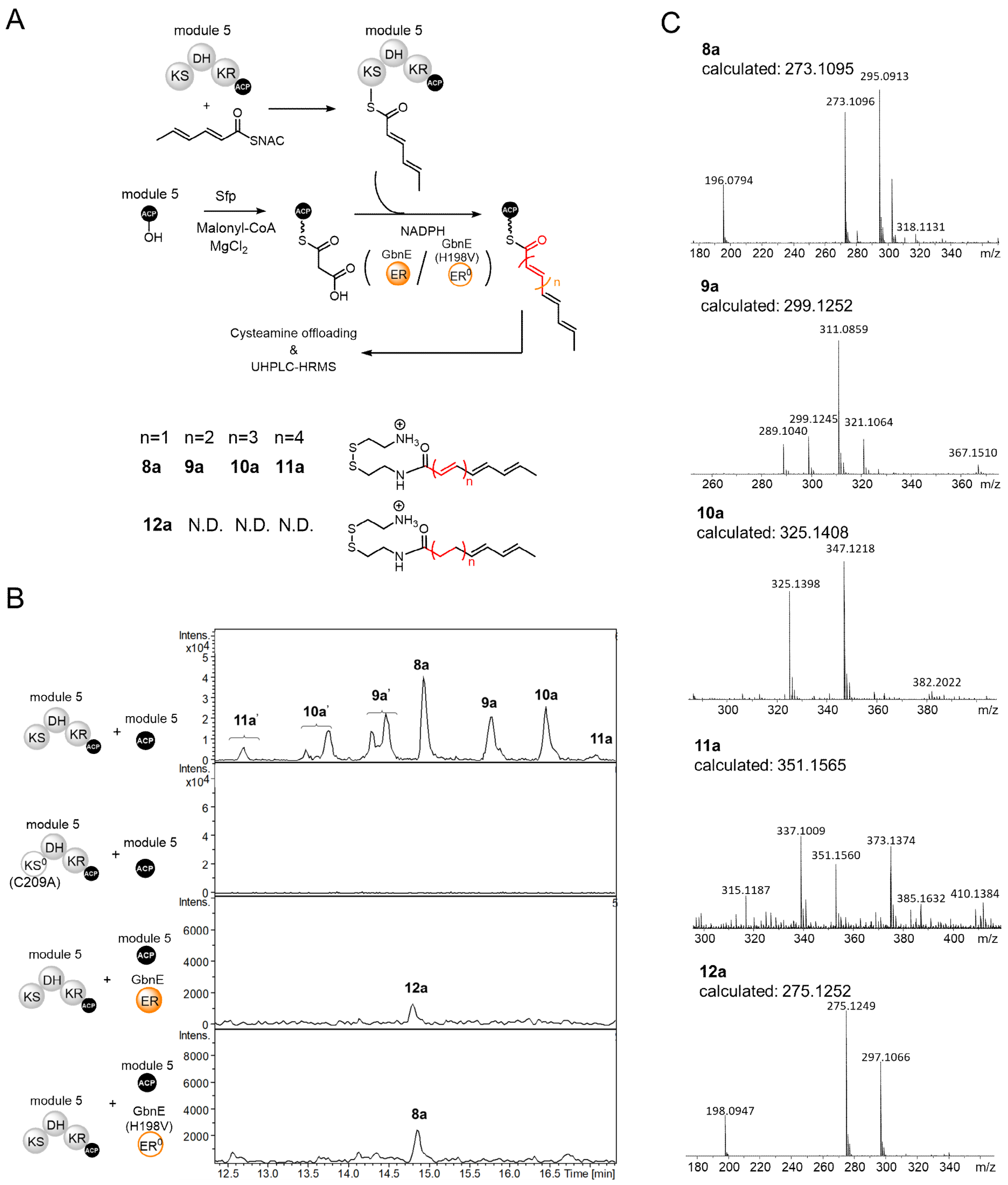


**Figure S8. UHPLC-ESI-Q-TOF-MS analyses of the cysteamine cleavage adducts for module 5 *in vitro* reconstitution assays, in the presence of the excised module 5 *holo*-ACP domain and GbnE or GbnE(H198V).** (A) Scheme illustrating the assay design. (B) Extracted ion chromatograms corresponding to the *m/z* values for the $[M+H]^+$ ions of cysteamine cleavage adducts **8a**, **9a**, **10a**, **11a** and **12a** resulting from reconstitution reactions in the presence and absence of GbnE, and the presence of the H198V mutant of GbnE. No products are observed in a control reaction lacking GbnE and employing a C209A mutant of module 5, which has a catalytically inactive KS domain. **9a'**, **10a'** and **11a'** are hypothesized to be the stereoisomers of **8a, 9a, 10a** and **11a** resulting from *E* to *Z* configurational isomerization of one or more double bond in the polyene. (C) High resolution mass spectra of **8a**, **9a**, **10a**, **11a** and **12a** and comparison with calculated *m/z* values for the $[M+H]^+$ ions.

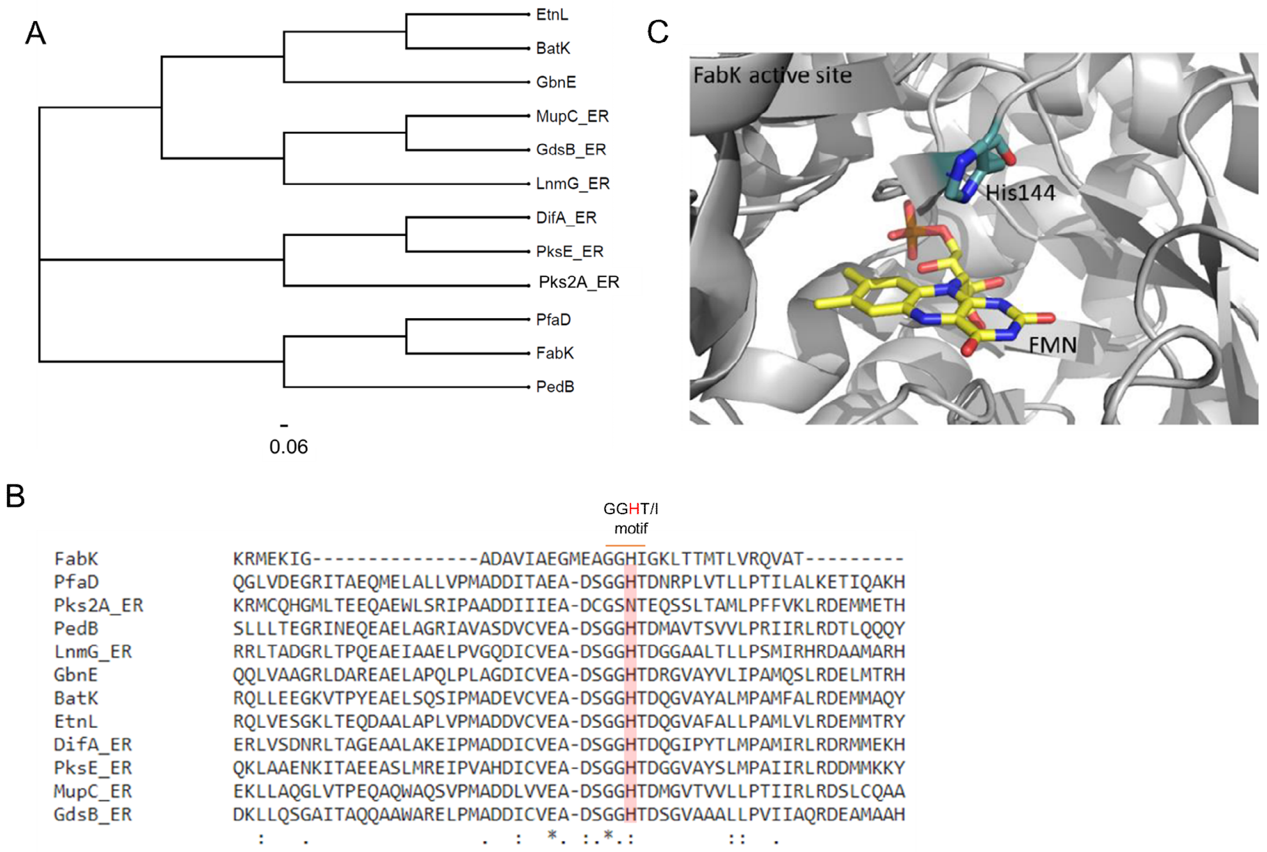


**Figure S9. Phylogenetic and sequence comparisons of *trans*-acting ERs from *trans*-AT PKSs with ERs from a polyunsaturated FAS and a type II FAS.** (A) Phylogenetic tree illustrating the evolutionary relationship between *trans*-acting ERs from *trans*-AT PKSs (Pks2A_ER: macrolactin[9], PedB: Onnamide[10], LnmG_ER: leinamycin[11], GbnE: gladiolin[12], BatK: batumin/kalimatacin[13], EtnL: etnangien[14], DifA_ER: Dificidin[15], PksE_ER: bacillaene/dihydrobacillaene[16], MupC_ER: mupirocin[17], GdsB_ER: gladiostatin[3]), PfaD (*trans*-acting ER from the type I iterative polyunsaturated fatty acid synthase in *Shewanella oneidensis* MR-1[8]) and FabK (ER from type II FAS in *Streptococcus pneumoniae*). (B) Region of a sequence alignment of the proteins used to construct the phylogenetic trees encompassing the conserved GGHT/I motif. A proposed catalytic residue (His144) in FabK and the corresponding residue in the other proteins are highlighted in red. (C) Structure of FabK active site (PDB ID: 2Z6I) showing the proximity of His144 to the FMN cofactor.

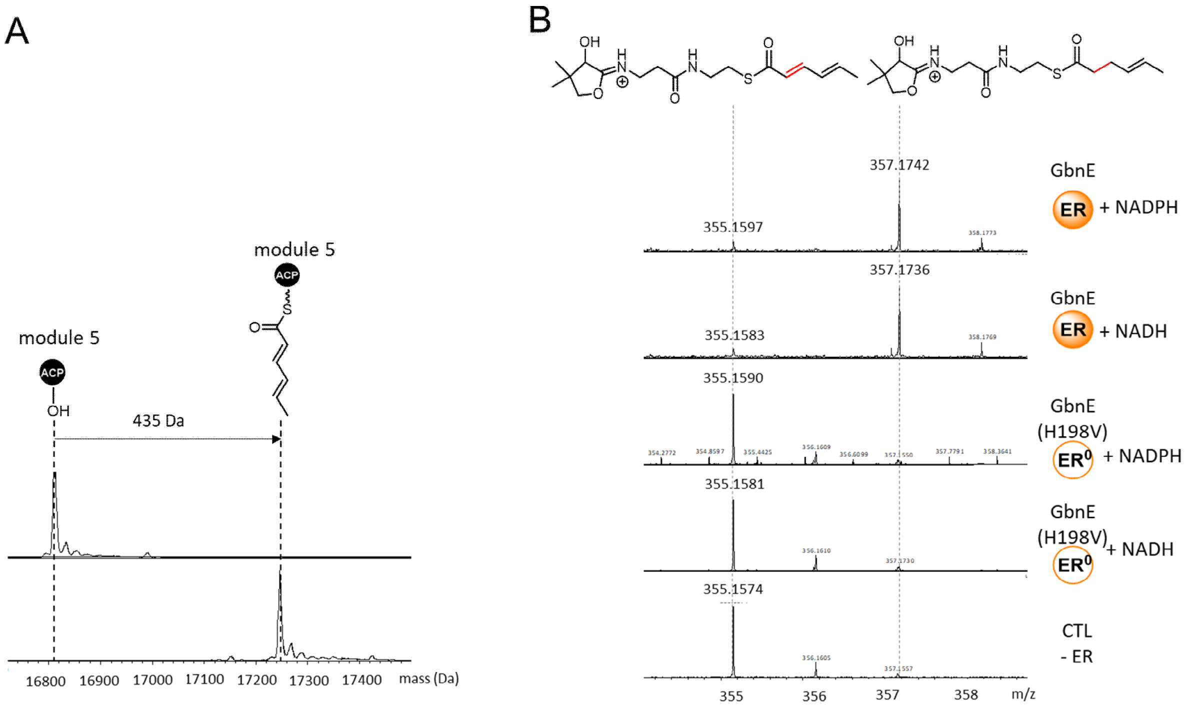


**Figure S10. *In vitro* characterization of GbnE(H198V).** (A) Deconvoluted mass spectra from UHPLC-ESI-Q-TOF-MS analyses of the excised module 5 *apo*-ACP domain from the gladiolin PKS before (top) and after (bottom) incubation with 2,4-hexadienoyl pantetheine thioester **17**, CoaA, CoaD, CoaE, sfp, $MgCl_2$ and ATP. (B) Ppant ejection ions observed for the hexadienoylated *holo*-ACP domain following incubation with GbnE and NADPH (top), GbnE and NADH (2nd from top), GbnE(H198V) and NADPH (3rd from top), GbnE(H198V) and NADH (2nd from bottom) and no ER (bottom) confirming GbnE(H198V) is catalytically inactive.

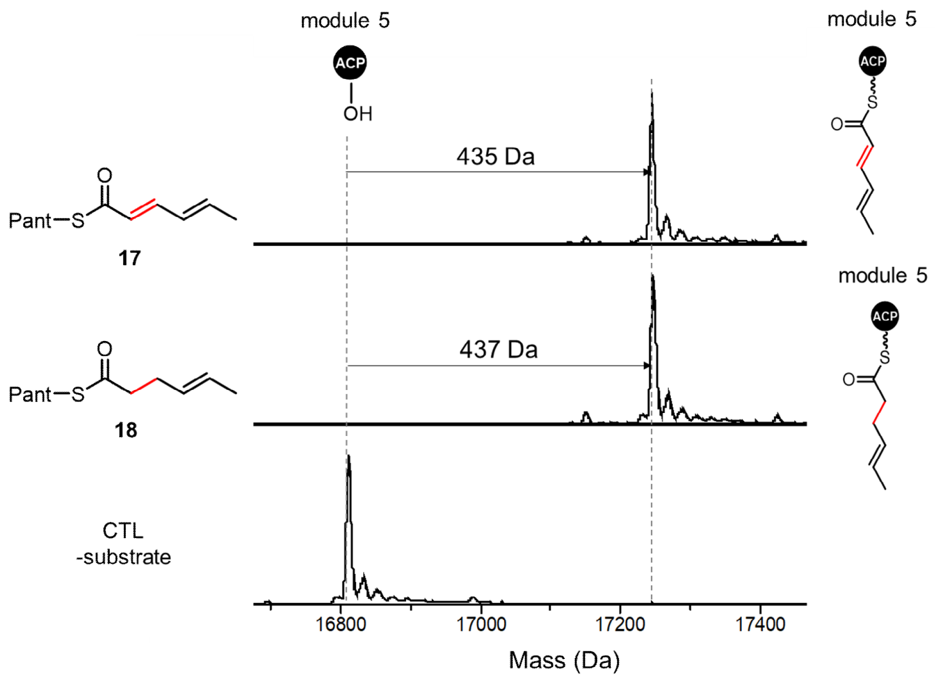


**Figure S11. UHPLC-ESI-Q-TOF-MS analysis of the conversion of the *apo*-ACP domain from module 5 of the gladiolin PKS to the 2,4-hexadienoylated- 4-hexaenoylated *holo* proteins**. Deconvoluted mass spectra of the *apo*-ACP domain after incubation with CoaC, CoaD, Sfp, $MgCl_2$, ATP and 2,4-hexadienoyl pantetheine thioester **17** (top), or 4-hexaenoyl pantetheine thioester **18** (middle), or no substrate (bottom).

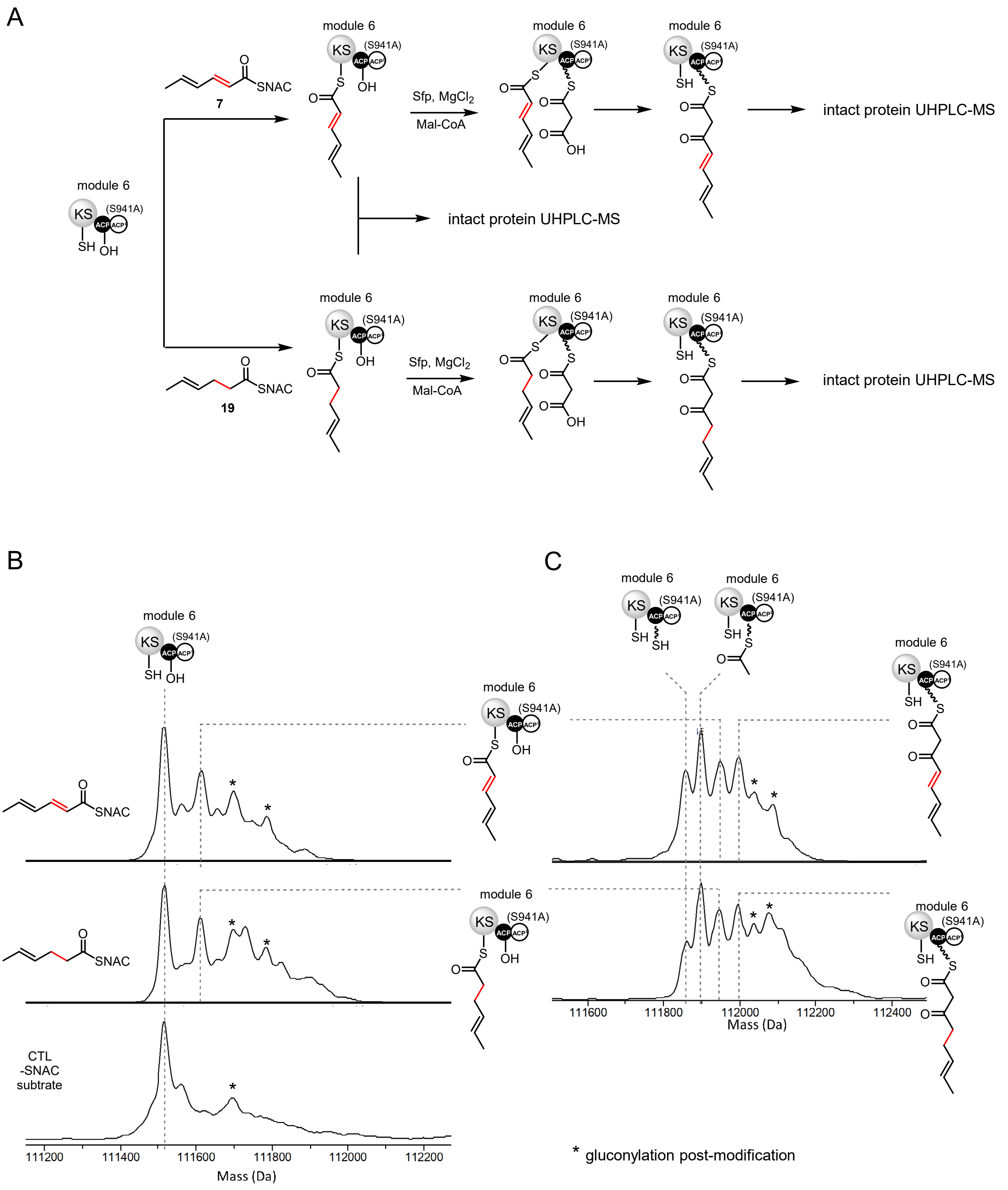


**Figure S12. *In vitro* characterization of the acyl donor substrate specificity for the chain elongation reaction catalyzed by KS domain from the module 6 of the gladiolin PKS**. (A) Scheme illustrating the workflow of the assay. (B) Deconvoluted mass spectra from UHPLC-ESI-Q-TOF-MS analysis of module 6(S941A) after incubation with 2,4-hexadienoyl NAC thioester **7** (top), 4-hexaenoyl NAC thioester **19** (middle), and no substrate (bottom). (C) Deconvoluted mass spectra from UHPLC-ESI-Q-TOF-MS analysis of module 6(S941A), acylated with the 2,4-hexadienoyl (top) or 4-hexaenoyl (bottom) unit, after incubation with Sfp, $MgCl_2$ and Malonyl-CoA. The asterisks denote gluconylated forms of proteins resulting from recombinant overproduction in *E. coli*.

### Codon optimized DNA sequences

##### >*etnL*

ATGATTACCGCCAAAAGTCTGGGTGACCGCAGTTTCAAAGCCGACTATGGCGTGGATTACGCCTATGTGGCGGGTGCGATGTATAAA
GGCATCGCGAGCAAGGAGCTGGTTGTTGCCATGGGCAAAGCCGGTTTTATCGCCTATCTGGGTACCGGTGGTCTGGATGAACGCGA
AATCGAGGCGAGCATTCGCAGCATTCAGAGCGTTATCGCGGGCCGTGCGTACGGCATGAATCTGCTGAGCAATCTGGAAAGCCCGG
AACTGGAGGAGCGTACCATCGATCTGTATCTGCGTCACGGTGTTCGTTGTGTTGAGGCCGCCGCGTATATGCGCGTTACGCCAGCG
CTGGTGCGTTATCGCCTCCAAGGTCTGGCCAGTGGCGCGGGTCGCACGCTGGTTGCCCCACGTCGTGTTCTGGCGAAAGTTAGTC
GCCCGGAAGTTGCGGCCGCGTTTATGCAGCCAGCCCCGGAAGCGGTGGTTCGTCAGCTGGTTGAGAGTGGCAAACTGACCGAAC
AAGATGCCGCGCTGGCCCCACTGGTGCCAATGGCCGATGATGTGTGCGTGGAAGCCGACAGTGGTGGCCATACCGATCAAGGTGT
GGCCTTCGCGCTGCTGCCAGCCATGCTGGTTCTGCGCGACGAGATGATGACGCGCTACCGTTACGAGAAACGCATCCGCGTTGGT
GCGGCGGGTGGTATTGGTACCCCACACGCGGCCGCCGCCGCCTTTGTTATGGGTGCGGATTTCATTCTGACGGGCAGCATTAACCA
GTGCACCCGTGAGGCCGGTACGAGCGAACCAGTGAAAGCGCTGCTGCAGCATCTGAACGTTCAAGATACCACCTACGCCCCGGCC
GGTGACATGTTCGAACTGGGTAGCAAGATCCAAGTTGTTCGCCGTGGTCTGTTCTTTCCAGCCCGCGCCAACAAACTGCACGAACT
CTACATGCGCCACGGCAGCCTCGAAGAAATTGACGCGAAAACGCGCCAGCAGATTCAAGAAAAGTACTTTCGCCGCAGCTTCGATG
CGGTTTGGAGCGAAACCCGCAGCTACTATGCGCGTACGTACCCGCATCGTCTGGCCGAGATCGAACGCAGCCCGAAACAGAAAATG
GCGGCCGTGTTCCGTTGGTACTTTGCGCATACCACCCGTCTGGCGCTGGAAGGTGTTGAAGACCAGCGCCTCGACTACCAGATCCA
TTGTGGTCCAGCGCTGGGCGCGTTCAATCAGTGGACCAAGGGTACGGCCATTGAACCATGGCAGAATCGCTACGTGGCCGACATC
GCCCGCCGTATTATGGAGGGTACCGCGGATCTGCTGAACGCCCGCTTTGGCGCCATGAAGGAAGCGGCGGAACCGCCACAGAGC
GTTAGCAGTTAA

##### >region encoding Etn_module 5 ACP domain

AGTCCCGGTGATCTGGCGGCCGCCAGTCCAGCGAATGCCAGCACCCCAGCCATTCGCGAAGAAGCGCTGCGCCAAGTTAAAAAAT
TATTAGCGCCGGTGATCAAACTGCCGGTGGAACGCATCGATGCCGCCGCCAGCTTTGAAACCTACGGTATCGAAAGCGTGATGGCC
GTTGAACTGACCGATCGTTTAGAAGCGGTGTTCGGTCCGCTGAGCAAAACGCTGCTGTTCGAAGCGAAAACCGTGCGCGAACTGG
CCGATTACTTTGTGGAACATCATGCCGCCACTTTAGGCTCTTTACTGGGCGGTGCGACCGCCCCGGCCGCCCCGGCCACCGCCGT
GAGCGCC

### Recombinant protein sequences

#### N-His$_6$-module 1 ER domain

```
MHHHHHHGKP IPNPLLGLDS TENLYFQGID PFT-
        10         20         30         40         50         60
AAPSPSGEAS TPLPAGDYGL VVRTVHALDE LSVEPWTPGE PGDDEVLIEV RASALNFPDV
        70         80         90        100        110        120
MCVQGLYPTQ PAYPFVPGFE VAGVVAAVGR AVVGIRVGEA VLALTGERMG GLASRVVVPA
       130        140        150        160        170        180
ANVLPKPSRL SFEEAASLPV AFLTAHHAFE TGRLAAGERV LIQTATGGCG LAAIQLARLR
       190        200        210        220        230        240
GARVYGTSSR AAKRALLERI GVEHVLDYRA AFDRELAGLT DGRGVDVVLN MLSGDAIQRG
       250        260        270        280        290        300
LDSLAPAGRY VEIAVHGLRT SGTLDLSRLV DNQSFHSIDL RRARARGLDW GETLAGLLPL
       310        320        330        340        350        360
FETGALVPIV SRVYPAEQLG EALRYVATGE HVGKVVIRHR GGALDDCVEA CVSRLVAQRE
       370        380        390
IAAREAARPA SLPVVPRRAS AAAQPASETH AP
```

#### N-His$_6$-module 1 ACP-ER didomain

```
MHHHHHHGKP IPNPLLGLDS TENLYFQGID PFT-
        10         20         30         40         50         60
PEPAEAGNVA VQATDADSFD LNISHDEIIE RSVAPDAATS RRRTIDSGPD LILERELAES
        70         80         90        100        110        120
LGQSLYLPAD EIDPERPFVE LGLDSIVGVE WARAINKAYG IALPATRLYD HVTIRLLAAH
       130        140        150        160        170        180
LVEQWGVARR DSARPAEPPV GLVAPVTRAP IAAAPSPSGE ASTPLPAGDY GLVVRTVHAL
       190        200        210        220        230        240
DELSVEPWTP GEPGDDEVLI EVRASALNFP DVMCVQGLYP TQPAYPFVPG FEVAGVVAAV
       250        260        270        280        290        300
GRAVVGIRVG EAVLALTGER MGGLASRVVV PAANVLPKPS RLSFEEAASL PVAFLTAHHA
       310        320        330        340        350        360
FETGRLAAGE RVLIQTATGG CGLAAIQLAR LRGARVYGTS SRAAKRALLE RIGVEHVLDY
       370        380        390        400        410        420
RAAFDRELAG LTDGRGVDVV LNMLSGDAIQ RGLDSLAPAG RYVEIAVHGL RTSGTLDLSR
       430        440        450        460        470        480
LVDNQSFHSI DLRRARARGL DWGETLAGLL PLFETGALVP IVSRVYPAEQ LGEALRYVAT
       490        500        510        520        530        540
GEHVGKVVIR HRGGALDDCV EACVSRLVAQ REIAAREAAR PASLPVVPRR ASAAAQPASE

THAP
```

N-His$_6$-SUMO-GbnE

MGSSHHHHHH GSGLVPRGSA SMSDSEVNQE AKPEVKPEVK PETHINLKVS DGSSEIFFKI
KKTTPLRRLM EAFAKRQGKE MDSLRFLYDG IRIQADQTPE DLDMEDNDII EAHREQIGG-

10 20 30 40 50 60
MAMITASSLG NADFRNDYRI KYAYLAGAMY RAIASKELVV ALGKAGLMGF LGTGGLRLDD
70 80 90 100 110 120
IEAAILQIQA ELGVDGVYGM NLLADLERPE REEATVDLYL RHGVRHVEAA AYMRVTPALV
130 140 150 160 170 180
RYRLRGASIG ADGRAVARHH VVAKVSRPEV ASQFMQPAPP ALVQQLVAAG RLDAREAELA
190 200 210 220 230 240
PQLPLAGDIC VEADSGGHTD RGVAYVLIPA MQSLRDELMT RHRYAKRIRI GAAGGIGTPQ
250 260 270 280 290 300
AAAAAFVMGA DFIVTGSINQ CTREAGTSDV VKALLQELDV QDTAYAPAGD MFELGAKVQV
310 320 330 340 350 360
ARRGLFFAAR ASRLHELYQQ HASLDEIDPA MRSQIEQKFF RRGFDEIWDE TRRHYASSRP
370 380 390 400 410 420
DQIAEIERSP KKKMAAVFRW YFAHSTQLAL SGDDTRLTDF QVHCGPALGA FNRWVRGTPL
430 440 450 460
ESWQNRHVAD LAERIMQGTA AWLEARLASM ADSGGQRPSL QLQRQ

N-His$_6$-module 3 ACP domain

MGSSHHHHHH SSGLVPRGSHM-

10 20 30 40 50 60
TATVPTTVTT PTATENDALA ARAIEYFRKW LAAQLKVGAD QLDDDAPLDR YGIDSVRVMQ
70 80 90 100 110 120
LTSALESRFG PLSKTLFFEY RNVAELSRHF VQAHRERMLD LLGLASPSAA PLPTSRPAGP
130
APRPVLSSGP AI

N-His$_6$-moudel 5 ACP domain

MHHHHHHGKP IPNPLLGLDS TENLYFQGID PFT-

10 20 30 40 50 60
PNPASLSSVP GGQDATRAAS AALLADEALR HVKRQLAAVI RLPVERIDED ASFEEYGIDS
70 80 90 100 110 120
VMAVELTDRL ERACGPLSKT LLFEYQSVRD LTAFLVRHHA EGLGAALGLG EASSDGETAK
AEV

N-His$_6$-moduel 10 ACP domain

MGSSHHHHHH SSGLVPRGSHM-

```
        10         20         30         40         50         60
AQAKPVAGAA PAAAGVPAPA PIADAAELGA RVDAALARAV CEILRIGESD VDAETDFSAY
        70         80         90        100        110        120
GFDSISLTEF SNRIGERLGV ELLPTIFYEY PSLLALRGFL LAEHGAALAA ALQVQPAAGR
       130        140
QVEPDPEPHR ESNRESNPGA AAVGLPEG
```

N-His6-moudel 12 ACP domain

MHHHHHHGKP IPNPLLGLDS TENLYFQGID PFT-

```
        10         20         30         40         50         60
VAAGYDGARA AALAAGESTR ASFDEALRRF VTDQLAAQGV ALAGRLGDDT PFFDAGLDST
        70         80         90        100        110
HLLALVRALE THCGRTFYPT LLFEHQTLRE LAAHLHRETP AAFGQAVPVW SESVAA
```

N-His6-module 5

MGSSHHHHHH SSGLVPRGSHMAS-

```
        10         20         30         40         50         60
AGARTMRAMA VATRGLAVPP ATASASLPQD RPEATATGTQ AVAVIGLAGR YPQAADLDAF
        70         80         90        100        110        120
WENLSTGRDC ITEIPSTRWD HEVYFDARKG QPGKSYSKWG GFLDGVDEFD PSFFSISPRE
       130        140        150        160        170        180
AQLMDPQERL FLQCAYHALE DAGHTRASLG AARVGVFVGV MYEEYPYHGS PSQGTTQPQA
       190        200        210        220        230        240
LGGSSASIAN RVSYAFNLNG PSIAMDTMCS SSLTALHLAC QSLRLGECEL ALAGGVNVSI
       250        260        270        280        290        300
HPNKYLGLSQ GQFASSEGRC RSFGAGGDGY VPSEGVGCVL LRPLAAAEAA GDRILGVIRA
       310        320        330        340        350        360
SAINHGGRTN GYTVPNPNAQ GELIAEALRA SGVDARAISY LEAHGTGTAL GDPIEIAGLV
       370        380        390        400        410        420
KAYGAWEGEP GEPGEPGDAR LEPCAIGSVK SNIGHCESAA GIAGLTKVLL QMRHGKLAPS
       430        440        450        460        470        480
LHAQTLNPLI DFGRTPFRVQ RELAPWRRPR VRVDGVEREM PRLAGISSFG AGGANAHVIV

       490        500        510        520        530        540
EEYVARAVEA ADARREGQPA IVVLSARSEA QVLIQARNLQ AAIEREAYGE GELAALAHTL
       550        560        570        580        590        600
QAGREAFEVR LATTVTSMAM LVERLASLAG EAPDYSAWMR GETRRDAADL PAPETVEGWL
       610        620        630        640        650        660
AQGRVEPLLR AWLDGLAFDW RRLRAADPPI RPLRLPGYPF ARQRYWADPP AAAAPRVARL
```

```
       670        680        690        700        710        720
HPLLHENRST AFEPQFVSHF DGREACLADH RVAHARVLPG AAQLEIARAA AALSLGEAAA
       730        740        750        760        770        780
LELTEVAWLQ PACFDEQGGS LRIALFVDSE TEAEFDIGSL DGDTVYSQGR LRLREAASPV
       790        800        810        820        830        840
VLELASLRAA CGQVPLAPEA LYASFAAAGL QYGPAHRAIA ELRTGLDTAG RPQVLARLEV
       850        860        870        880        890        900
PDGAADEALV LHPSLVDGAI QATIGLLLVP GGARRAMLPA RLGRVGVAAA TPARGWAWLR
       910        920        930        940        950        960
FAEGSGPEDV APRIDLSICD EHGAVALEIE ALTLMPAPVA SQTGVETLWL APAWAEAEAP
       970        980        990       1000       1010       1020
LDAPRGAAQP TREVFVAGQL PEGVLAALAR RYGESRVHVA PGADAMPLAA GYVAAADALF
      1030       1040       1050       1060       1070       1080
AIVRERLADV ASGECLFQLL RVEDGTAGAA CLDGLGALLR TASIENPRFA TQSIRIDAAD
      1090       1100       1110       1120       1130       1140
ATDGETLLAL LDANARERDS REIAYLDGRR RQRRHLELSQ PGGDVPAWPA GTVALITGGL
      1150       1160       1170       1180       1190       1200
GGIGYRVAES IAASGPGTTL LLCGRSEPAD AAARLGALRA AGANAEFVRT DITDAAATEA
      1210       1220       1230       1240       1250       1260
LVAGIVARHG RLDTVIHSAG IVRDNFVIRK NASELHAVLA PKVAGLVNLD AATRSLPLRE
      1270       1280       1290       1300       1310       1320
LIVFSSVSGA FGNAGQADYA CANAFMDAFA AWRATRVAAG ERSGRTLSIG WPLWAEGGMR
      1330       1340       1350       1360       1370       1380
VDAAVEAKLR RDGLAPLDTA SGLAVLARCR QLPPELTQVV VLAGDAARLR ARHGLAAAAI
      1390       1400       1410       1420       1430       1440
GAAASAPNRA PNPAPNPASL SSVPGGQDAT RAASAALLAD EALRHVKRQL AAVIRLPVER
      1450       1460       1470       1480       1490       1500
IDEDASFEEY GIDSVMAVEL TDRLERACGP LSKTLLFEYQ SVRDLTAFLV RHHAEGLGAA
      1510
LGLGEASSDG ETAKAEV
```

N-His$_6$-module 6

```
MGSSHHHHHH SSGLVPRGSHMAS-
        10         20         30         40         50         60
DGETAKAEVA VAALAAGAVA TARAAGPALA PVAPRTRRRP RGRLPGRESE PPRITAIAVI
        70         80         90        100        110        120
GLAGRYPQAA DLDAFWENLS TGRDCITEIP STRWDHEAYF DARKGQPGKS YSKWGGFLDG
       130        140        150        160        170        180
VDEFDPFFFN ISPREAQLMD PQERLFLQCA YHALEDAGHT RASLGAVRVG VFVGVMYEEY
       190        200        210        220        230        240
HLLSDPAGGD LAQTYIPGGY LSSVANRVSY FGNFRGPSFG VDTMCSSSLT ALHLACQSLR
       250        260        270        280        290        300
LGECELALAG GVNVSIHPNK YLGLSQGQFA SSEGRCRSFG AGGDGYVPSE GVGCVLLRPL
```

```
       310        320        330        340        350        360
AAAEAAGDRI LGVIRASAIN HGGRTNGYTV PNPNAQGELI AEALRASGVD ARAISYLEAH
       370        380        390        400        410        420
GTGTALGDPI EIAGLVKAYG AWEGEPGEPG DARLEPCAIG SVKSNIGHCE SAAGIAGLTK
       430        440        450        460        470        480
VLLQMRHGKL APSLHAQTLN PLIDFGRTPF RVQRELAPWR RPRVRVDGVE REMPRLAGIS
       490        500        510        520        530        540
SFGAGGANAH LIVEEYVARA VEAADARREG QPAIVVLSAR SEAQVLIQAR NLQAAIAREA
       550        560        570        580        590        600
YGEGELAALA HTLQAGREAF EVRLATTVTS MAMLVERLAS LAGETPDYSA WMRGEARRDG
       610        620        630        640        650        660
NDALAHFAKD PELNEVLRKW LRAGNPVRIA ELWTRGLDID WAAMQEDGPA PARLRLPGYP
       670        680        690        700        710        720
FATKRYWMAR ADGSPALAAR AAVPARPEPP TRAAVPPAAV KVLAEVLADA PRVAIVLGEP
       730        740        750        760        770        780
GGFEARTEGA ARAASIRLAV LDPLDEFEPP AAPAAARQDD GQTAAAARQV PAGIEVSIER
       790        800        810        820        830        840
LRLSLAQMLC AEAADLGADV PFGELGLDSV VGVEWTRAIG REYGVTLAAA TLYDHPTLTT
       850        860        870        880        890        900
LAAWLAATVE AGAAIATAGR DVPPRPDAHP VRLVEPDQDV ATPLAAAAAV EGGNEAEVAA
       910        920        930        940        950        960
GMAMPIEALV ERLAGTLAQA LCAEREEIDP DTPFAELGID SVVGVEWVRD INRAFGTELK
       970        980        990       1000       1010       1020
ATVLFDHATV KLLAAHLAPL ACPAPHSVRV DPPAAPVEQL AAASAVPARE LADTARHAGS
      1030       1040
RGHVQAVEPA APVAKTGNAP
```

N-His6-MBP-EtnL

MGSSHHHHHHSSGLVPRGSHMASGKIEEGKLVIWINGDKGYNGLAEVGKKFEKDTGIKVTVEHPDKLEEKFPQVAATGDG

PDIIFWAHDRFGGYAQSGLLAEITPDKAFQDKLYPFTWDAVRYNGKLIAYPIAVEALSLIYNKDLLPNPPKTWEEIPALD

KELKAKGKSALMFNLQEPYFTWPLIAADGGYAFKYENGKYDIKDVGVDNAGAKAGLTFLVDLIKNKHMNADTDYSIAEAA

FNKGETAMTINGPWAWSNIDTSKVNYGVTVLPTFKGQPSKPFVGVLSAGINAASPNKELAKEFLENYLLTDEGLEAVNKD

KPLGAVALKSYEEELAKDPRIAATMENAQKGEIMPNIPQMSAFWYAVRTAVINAASGRQTVDEALKDAQTRITKGENLYF

QGS-

```
        10         20         30         40         50         60
MITAKSLGDR SFKADYGVDY AYVAGAMYKG IASKELVVAM GKAGFIAYLG TGGLDEREIE
        70         80         90        100        110        120
ASIRSIQSVI AGRAYGMNLL SNLESPELEE RTIDLYLRHG VRCVEAAAYM RVTPALVRYR
       130        140        150        160        170        180
LQGLASGAGR TLVAPRRVLA KVSRPEVAAA FMQPAPEAVV RQLVESGKLT EQDAALAPLV
```

```
        190         200         210         220         230         240
PMADDVCVEA  DSGGHTDQGV  AFALLPAMLV  LRDEMMTRYR  YEKRIRVGAA  GGIGTPHAAA
        250         260         270         280         290         300
AAFVMGADFI  LTGSINQCTR  EAGTSEPVKA  LLQHLNVQDT  TYAPAGDMFE  LGSKIQVVRR
        310         320         330         340         350         360
GLFFPARANK  LHELYMRHGS  LEEIDAKTRQ  QIQEKYFRRS  FDAVWSETRS  YYARTYPHRL
        370         380         390         400         410         420
AEIERSPKQK  MAAVFRWYFA  HTTRLALEGV  EDQRLDYQIH  CGPALGAFNQ  WTKGTAIEPW
        430         440         450
QNRYVADIAR  RIMEGTADLL  NARFGAMKEA  AEPPQSVSS
```

N-His$_6$-Etn module 5 ACP domain

```
MGSSHHHHHHSSGLVPRGSHM-
         10          20          30          40          50          60
SPGDLAAASP  ANASTPAIRE  EALRQVKKLL  APVIKLPVER  IDAAASFETY  GIESVMAVEL
         70          80          90         100         110
TDRLEAVFGP  LSKTLLFEAK  TVRELADYFV  EHHAATLGSL  LGGATAPAAP  ATAVSA
```

N-His$_6$-Etn module 10_ACP domain

```
MGSSHHHHHHSSGLVPRGSH-
         10          20          30          40          50          60
TPASEQPAPL  ADGAGEASGL  RDLVERTLVR  AVSRVLKIHE  ADIELDAELS  AFGFDSLSLT
         70          80          90         100         110
ELSNRLSAEL  GVELLPTVFF  EHPSLAALET  FLLATHLGAL  ARTFRAEVAA
```

N-His$_8$-EtnK_AT

```
MKHHHHHHHH  GGLVPRGSHGSH-
        10          20          30          40          50          60
AGPRATVAFM  FPGQGSQKRG  MGAGLFDSVP  EYAAVEREVD  ALLGYSMRAL  CHEDPDGRLK

         70          80          90         100         110         120
ETQYTQPALY  VVNALHYYDA  IARGQRPDYV  AGHSLGEYNA  LLAAGAFDLL  TGLRLVKKRG

        130         140         150         160         170         180
ELMAASVSGG  MAAVIGMDEG  RIKQVLTENG  LGTIDVANFN  SPSQIVISGP  VADIARGHTV

        190         200         210         220         230         240
FEDAGARTYV  ILPVSAAFHS  RYVEETGRAF  ADFIAPMRFE  ALRIPVISNV  SARPYEAKDP

        250         260         270         280         290
SATIKSLLVQ  QITRPVQWVQ  SVSFLMAQGV  KEFREIGPGN  VLTRLVQQIE  RQPSGA
```

(10) Piel, J.; Hui, D.; Wen, G.; Butzke, D.; Platzer, M.; Fusetani, N.; Matsunaga, S. Antitumor polyketide biosynthesis by an uncultivated bacterial symbiont of the marine sponge *Theonella swinhoei*. *Proc. Natl.*

*Acad. Sci. U. S. A.* **2004**, *101* (46), 16222–16227.
